# Distinct interferon response patterns link to increased transposable element expression in leukocytes of systemic lupus erythematosus patients

**DOI:** 10.1101/2025.05.30.657099

**Authors:** Leo J. Arteaga-Vazquez, Hugo Sepulveda, Bruno Villalobos Reveles, Kazumasa Suzuki, Kenneth C. Kalunian, Ferhat Ay, Mark R. Boothby, Anjana Rao

## Abstract

**Background:** Systemic lupus erythematosus (SLE) is a spontaneous systemic auto-immune condition for which the inciting factors and genetic basis are generally unknown. Although heterogeneous in its manifestations and severity, SLE involves chronic inflammation along with sustained autoantibody production. The root causes and pathophysiology of the inflammation and breaches of tolerance are incompletely understood, but neutrophils are thought to be important elements of the pathophysiology. Type I interferons (IFN) in the bloodstream and an IFN-stimulated gene (ISG) signature in circulating leukocytes, including neutrophils, are common features in many patients. Earlier work has provided evidence of increased levels of transcripts derived from transposable elements (TEs) in peripheral blood cells of SLE patients. Using six leukocyte types, including neutrophils, we tested the correlation of TE expression with disease severity and explored the relationships between increased ISG and TE expression with attention to the genomic locations of the expressed TEs.

**Results:** We reanalysed previously published data from neutrophils and other leukocytes of SLE patients sub-divided into ISG-high (termed IFNpos, n=12) and ISG-low (termed IFNneg, n=11) patients in the original study, examining RNA-seq data from B and T lymphocytes, conventional and plasmacytoid dendritic cells (DC), monocytes and PMN of IFNpos and IFNneg SLE patients compared to healthy controls. SLE patients pre-stratified as IFNneg showed no significant increase in TE expression. All IFNpos cell types had similar amounts of total TE-encoded RNA, but among the 6 cell types, PMN had the highest number of differentially expressed TEs and ISGs in IFNpos SLE patients compared to healthy controls. There was a strong correlation between expression of several specific TE families and disease activity assessed at the time of the visit. Most upregulated TEs (∼80%) were present in introns of upregulated genes, and ∼67% of these were ISGs. By mapping expressed TEs in ISGs, we found that high intronic TE expression correlated strongly with increased ISG expression as well as with splicing alterations in annotated exons flanking expressed TEs. Consistent with autonomous TE expression, upregulated TEs were also observed at intergenic sites distant from annotated genes, perhaps due to weakening of heterochromatin integrity.

**Conclusions:** Our findings show a strong association and suggest mechanistic relationships between increased TE expression and IFN responses in multiple types of leukocytes centrally involved in SLE pathogenesis. Although limited by short-read RNA-seq technology, our analyses support selective upregulation of some TEs independent from the regulation of conventional genes, concurrent with many intron-localized TEs whose expression tracks with ISGs. The data emphasize the need for long-reads sequencing to understand the causes and consequences of high TE expression in SLE and other autoimmune/inflammatory disorders. Important questions include whether TE expression in introns of ISGs and other genes is independently regulated or reflects exonization or partial intron retention, and how frequently it correlates with splicing variations in adjacent exons.

## INTRODUCTION

Systemic lupus erythematosus (SLE) is a prototypical example of a chronic autoimmune disorder, a class of systemic conditions that involve persistent auto-immune dysfunction with inflammation [1–3]. The underlying causes, pathophysiological mechanisms, and optimal treatments for most of these chronic illnesses remain to be fully elucidated. The demography, clinical features, and apparent causes for SLE vary substantially [2–4]. Nonetheless, breaches of the tolerance mechanisms that ordinarily prevent chronic and sustained production of antibodies directed against “self” or “modified self” determinants [3–6] are a central feature of SLE. These auto-antibodies can drive inflammation in various tissues.

Polymorphonuclear granulocytes (PMN), also known as neutrophils, are a prominent pathogenic cell type in SLE. PMNs undergo cell death concomitantly with a process known as NETosis [7–9] – externalisation of a chromatin meshwork that releases double-stranded DNA (dsDNA) and intranuclear antigens that elicit auto-antibodies that are a characteristic feature of SLE. There is also a strong connection between SLE and interferon (IFN)-stimulated genes (ISG) expression. Many SLE patients display a gene expression signature enriched for type I interferons (IFN-I, IFNα and IFNβ) and interferon-stimulated genes (ISGs) (reviewed in [10–14], and sera of subjects who later developed SLE had subtly higher levels of circulating IFN-I and IFN-γ as well as auto-antibodies even before the onset of clinical disease [15–17]. A variety of cell types can be sources of IFN-I, but PMNs have been suggested as a major contributor [18]. Recent studies highlight the correlation of ISG expression with increased TE expression [19], but the genomic locations of upregulated TEs and whether they influence, or are influenced by, conventional protein-coding gene expression is not clear. Moreover, pointing to the need for additional analyses, a recent study [20] that detected increased TE expression in leukocytes of autoimmune disease patients found no such increases in subjects with SLE. The mechanisms responsible for the initiation of increased IFN production in most SLE patients are not yet known, and indeed the root causes may differ among SLE patients. In some instances, however, altered regulation of chromosome silencing or inadequate control of immunostimulatory RNA have been proposed as important in the pathophysiology of SLE [21–25].

In eukaryotic cells in general, and metazoans in particular, health and longevity depend on a capacity to preserve the integrity of the nuclear and mitochondrial genomes along with the epigenetic networks that stabilize cellular identities while allowing changes in gene expression [26, 27]. One challenge to genome stability is posed by the invasion of genomes by transposable elements (TEs) over millions of years of evolutionary time [28–34]. After the initial invasion, TEs can expand their numbers significantly, creating large families and subfamilies of repeated TEs that include LTR-containing endogenous retroviral sequences (ERVs) as well as short and long interspersed nuclear elements (SINEs and LINEs, respectively) [28–34]. Over evolutionary time, these processes result in the generation of TEs that are nonfunctional, in part due to severe mutations in older elements, and in part due to the emergence, through evolution, of KRAB zinc finger protein (KRAB-ZFP) proteins that bind to the regulatory regions of some TEs and suppress their expression via their interaction with KAP1 [28, 33]. TEs and mutated TE-derived sequences are estimated to constitute ∼40-50% of the haploid genome in humans [29, 35].

Repetitive sequences, including TE families, are sensed as threats by the non-repetitive genome due to their potential for destabilizing genomic sequences by transposition and recombination [27, 33]. In consequence, TE expression – required for transposition – is suppressed by a variety of mechanisms, among them mutations, DNA methylation, heterochromatin-associated histone modifications such as H3K9me3, and KAP1-KRAB ZFP and HUSH protein complexes [28, 29, 33, 36, 37]. Errors in these mechanisms would lead to a slow loss of heterochromatin integrity – termed heterochromatin “weakening” or “dysfunction” – which could potentially contribute to disparate diseases such as aging and cancer [37–39]. A reliable marker for heterochromatin dysfunction is increased expression of transposable elements (primarily LINEs and LTR elements) that are normally silenced in heterochromatin [37–41].

When heterochromatin integrity is compromised by decreased DNA or H3K9 methylation, TE regulatory regions (LTRs of endogenous retroviruses and the 5’ untranslated regions (5’-UTR) of LINEs) can participate in the regulation of neighboring gene transcription by acting as distal enhancers or alternative promoters, or they can modify gene structures by becoming alternative exons [28, 30–32, 40–43]. Further, the repetitive nature of TE sequences creates a potential for recombination events, genomic stress and increased mutations, and TE expression correlates with the inflammation characteristic of cancer, aging and autoimmune disease. Specifically, micronuclei produced by chromosome mis-segregation can activate cGAS/STING pathways [44–46], and double-stranded RNA encoded by inverted-repeat Alu elements can activate innate sensor mechanisms, especially if ADAR-mediated RNA editing is compromised, culminating in MDA5 activation akin to an anti-viral response [47, 48]. The consequent release of type I interferons (IFN-I) and the increased transcription of interferon-stimulated genes (ISGs) can evoke systemic inflammation and autoimmune disease. In support of such a mechanism, GWAS studies of several autoimmune diseases have linked ADAR loss-of-function and MDA5 gain-of-function alleles to diverse autoimmune/ inflammatory conditions including SLE [49–51].

These findings raise questions about the molecular and cellular mechanisms that initiate or contribute to perpetuating the dysregulated production of type I interferons in SLE. A variety of cells can serve as sources of type I interferons when exposed to stimuli such as apoptotic cells, circulating nucleoprotein complexes derived in part from PMN, and oxidized mitochondrial DNA among others. However, as noted above, de-repression of TE transcripts may offer another potential bridge connecting innate sensor mechanisms to increases in IFN-I [52–58]. With these considerations in mind, we measured TE-derived RNA in PMN and several other types of leukocytes of SLE patients compared to healthy control (HC) individuals. To ascertain if TE de-repression is a general feature in SLE, and to gain insight into commonalities or differences among several WBC types, we examined the relationship between selected RNAs that could be mapped to specific TE locations in the genome and RNA from genes in their vicinity. We also tested the effect of stratifying SLE patients by whether they displayed or lacked a pre-defined interferon signature based on their expression of interferon-stimulated genes (ISGs). We then investigated the relationships between (i) the IFN signature and TE expression, (ii) TE expression and clinical disease, in individual SLE PMN samples stratified by whether they belonged to the ISG-high (IFNpos) or ISG-low/negative (IFNneg) subset as defined in prior work [59]. We found that such stratification increased the sensitivity of detection of differentially expressed TE families, relative to analyses in which all SLE patients were merged without regard to IFN signatures. Stratification into IFNpos and IFNneg subsets also revealed that patients with severe versus moderate or mild disease could be distinguished at least as well by quantification of the expression levels of these differentially expressed TEs as by their IFN signature. Finally, we localized upregulated TEs whose genomic origin could be uniquely mapped, and documented high frequencies of expressed TEs in introns, frequently but not exclusively those of upregulated ISGs.

## RESULTS

### TEs show increased expression in PMN and other leukocytes from IFNpos but not IFNneg SLE patients relative to healthy controls

In a majority of SLE patients, the peripheral blood leukocyte pool exhibits increased levels of ISGs [10–14]. A prior analysis of RNA-seq data from SLE patients and healthy control donors identified two subgroups of SLE patients, with high (“IFNpos”) and low (“IFNneg”) ISGs expression, respectively [59]. In this study, polymorphonuclear leukocytes (PMNs, also known as neutrophils) in peripheral blood showed a strong upregulation of TNFSF13B (BAFF), a cytokine that supports B cell survival and can cause loss of self-tolerance when available to B cells in excess [60, 61]. Neutrophils are important mediators of SLE pathogenesis [7–9, 18] and have been reported to express LINE-1 RNAs and their encoded protein products [53]. Accordingly, we elected to perform in-depth re-analyses of the RNA-seq data from purified PMN reported in this paper [59], so as to explore more closely the TE dysregulation in SLE patients and any association with the severity of the disease.

We compared RNA-seq data from 10 healthy controls (HC) to 23 SLE patients classified by their ISG levels as IFNneg (n=11) and IFNpos (n=12) (**Fig 1A**, *top;* Supplemental Fig. 1A) [59]. The overall numbers of differentially expressed (DE) ISGs and TE subfamilies were highest in PMN among the leukocyte types originally analyzed (**Fig 1B**), prompting us to focus first on this cellular subset. To weight each subject equally, we restricted our analysis to only the initial visits (**Suppl. Fig. 1A**). In a Principal Components Analysis (PCA) of the RNA-seq datasets, the 12 IFNpos SLE samples were clearly distinguished from the 10 healthy control samples and from IFNneg samples (**Fig 1A**, *bottom*). The PCA analysis also showed that, despite clear diagnoses of SLE in all IFNneg patients, the main components of overall gene expression in IFNneg samples largely overlapped with those of the healthy controls. This finding is consistent with previous studies suggesting distinct subtypes among SLE patients, including a subset of patients in whom the IFN signature may be muted or absent [62]. Compared to algorithms used in other studies, the gene network analysis identified a higher fraction of the patients in the donor populations used here as IFNneg, consistent with the heterogeneity of IFN effects in the peripheral leukocytes of SLE patients. Using an extended list of ISGs (**Suppl. Table 1**) ([40, 41], modified from [48]), pairwise analyses of differential ISG expression between sample sets showed the expected difference between the pool of all 23 SLE patients and healthy controls (**Fig 1C**). The differences were of greater magnitude when IFNpos were compared with healthy controls (**Fig 1D, E**). In line with the PCA and the gene network stratification, ISG expression in IFNpos samples was of substantially greater magnitude than in IFNneg patients (**Fig 1F**).

**Figure 1.**
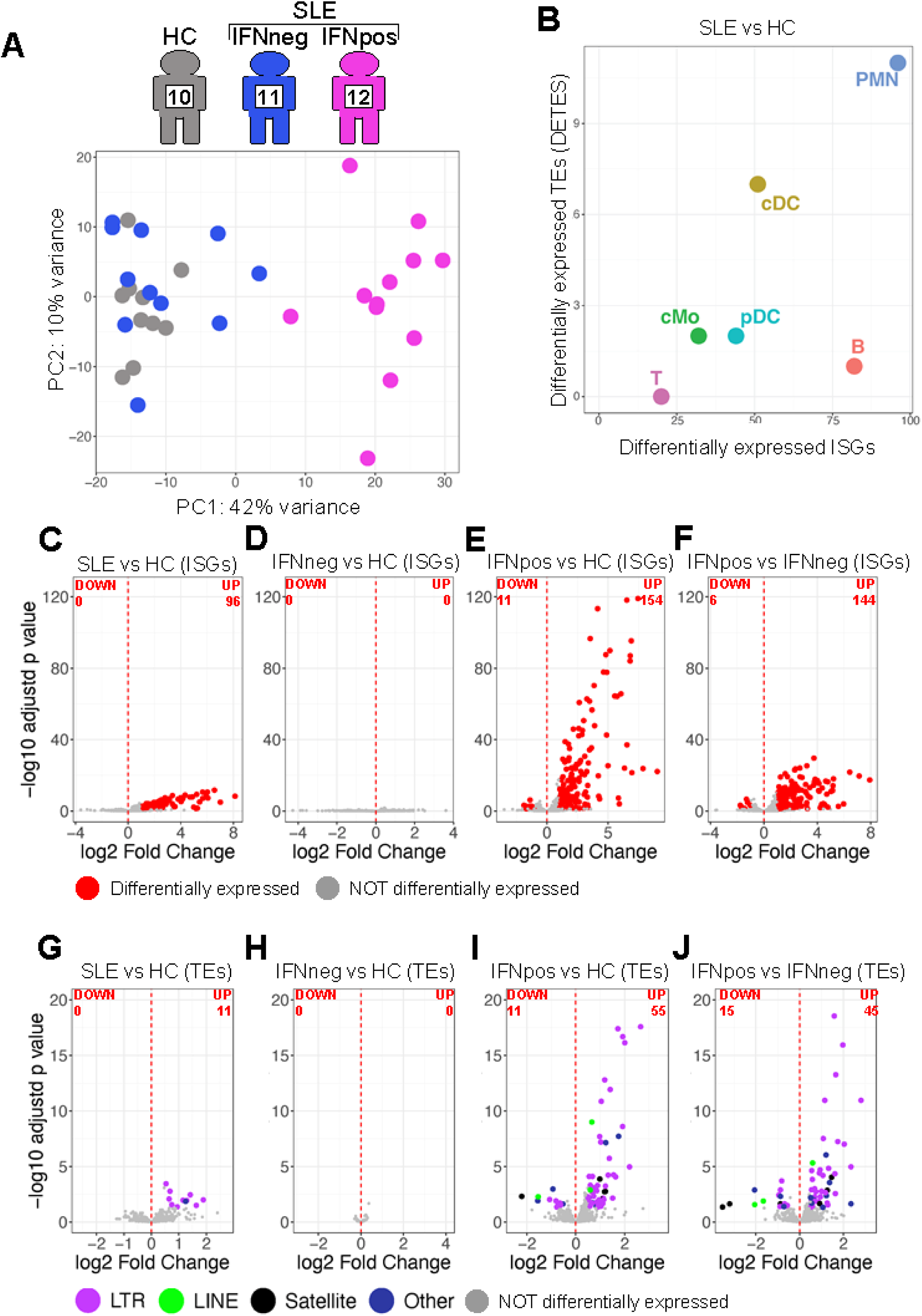
Increased TE expression in the PMN of SLE patients is confined to the IFNpos subset. A, *Top*, the sample numbers for patients in the subgroups (HC, healthy control; each subset of SLE). *Bottom*, A PCA plot of RNA-seq data for all PMN samples color-coded as in the homunculi (HC, gray; IFNneg, blue; IFNpos, mauve). **B,** A plot of differentially expressed TEs versus ISGs (P_adj_ ≤ 0.05) for all cell types purified from peripheral blood, using RNA-seq data from all 23 SLE patients and 10 healthy controls (HC). T, T cells; cMo, conventional monocytes; pDC, cDC, plasmacytoid and conventional DC; B, B cells; PMN, neutrophils). **C-E,** Volcano plots of pairwise comparisons between groups, using RNA-seq data from PMN and a list of ISGs (Supplemental Table 1). Specific comparisons: **C,** All 23 SLE vs HC; **D,** IFNneg SLE samples vs HC; **E,** IFNpos SLE samples vs HC; **F,** IFNpos vs IFNneg subsets of SLE. **G-J** As in C-E, except that TE sequences were analyzed, categorized and color-coded as indicated (LTR, LINE, Satellite, Other). **G,** All SLE vs HC; **H,** IFNneg SLE vs HC; **I,** IFNpos SLE vs HC. **J,** TEs in the comparison of the IFNpos to the IFNneg subset.

The literature suggests many connections between TE expression and inflammation in cancers, autoimmune diseases and aging (reviewed in [37, 39]), but the connections between TE and ISG expression are not yet understood. Applying the TEtranscripts pipeline (*see Methods*), which classifies and quantifies TE-related transcripts, showed the expression of several TE families and subfamilies to be increased significantly in SLE PMN (**Fig 1G**), with LTR (endogenous retrovirus, ERV) sequences dominating the observed differentially expressed (DE) TEs. No TEs were differentially expressed when comparing patient samples stratified as IFNneg to those of healthy controls (**Fig 1H**). In sharp contrast, the expression of multiple TE families and subfamilies was strongly increased in IFNpos SLE patients compared to healthy controls (**Fig 1I**) or to IFNneg SLE patients (**Fig 1J**). The stratification to IFNpos increased the sensitivity of detecting increases in TE-encoded RNA five-fold when compared to the total population of all SLE patients (i.e., 55 TEs upregulated in the IFNpos to HC comparison versus 11 TEs upregulated in the total SLE to HC comparison, **Suppl. Fig 1B**, compare panels 3 and 1); with all TE classes (LTR, Satellite, LINE, and DNA transposons) detected (**Fig 1I; Suppl. Fig. 1B**).

Examination of other cell types (B and T lymphocytes; conventional and plasmacytoid DC; conventional monocytes) showed that fewer DE TEs were found in any of the cell types, especially if comparing all SLE to HC, yet such upregulation was readily detected in each cell type when comparing IFNpos to HC or IFNneg samples (**Suppl. Fig 2**). The increased sensitivity afforded by this stratification allowed identification of multiple upregulated RNAs derived from LINE and SINE elements in addition to the majority of LTR-family TEs in most cell types other than PMN. SINE and DNA TEs constituted a larger fraction of upregulated DE TEs in classical monocytes (cMo), in contrast to PMN and the other leukocyte types. Hence RNA encoded by many classes and families of TEs are increased in PMN and other leukocytes of SLE patients, but in this moderate-sized and diverse patient pool, the increases are predominantly in cells from SLE patients classified as IFNpos.

### Relationships between ISG and TE expression in PMN of SLE patients

The IFNpos SLE samples showed transcriptional profiles distinct from those of IFNneg and HC samples (**Fig 1A**), and a congruent result was obtained when samples were clustered based on the expression of ISGs or TEs (**Fig 2A, B**, respectively). In these heatmaps with dendrograms, IFNpos SLE patients clearly differed from healthy control and IFNneg SLE samples; the IFNneg SLE and healthy control samples were comingled and more similar to one another. Gene set enrichment analyses (GSEA) showed enrichment of interferon response pathways in all SLE versus healthy control PMN, as well as in IFNpos versus healthy controls or IFNneg as expected (**Fig. 2C**). Although there was no general enrichment of differentially expressed ISGs in PMN of IFNneg SLE patients’ versus healthy controls (**Fig. 1D**), a slight enrichment of the interferon alpha response pathway was observed (**Fig 2C**). Comparing PMNs from healthy controls versus SLE patients, the numbers of differentially expressed total genes, ISGs and TEs were significantly higher in IFNpos vs HC (**Fig 2D**). These results show that the heterogeneity among SLE samples can be observed not only in the interferon signature, but also in TE expression.

**Figure 2.**
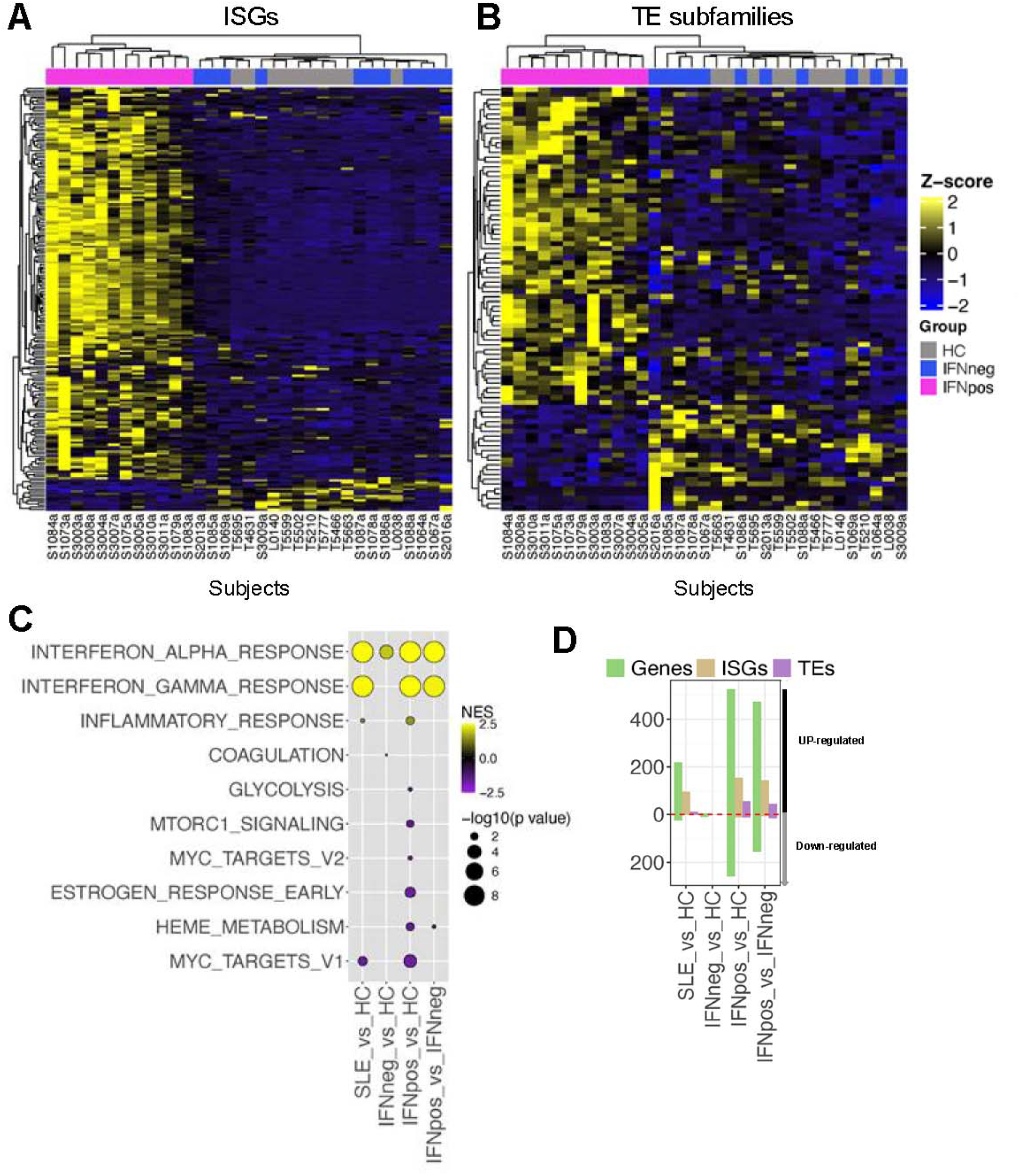
Distinct characteristics of the IFNneg and IFNpos subsets. A, A self-organizing heatmap of PMN samples using differentially expressed ISGs. Each row represents an individual gene of the ISG signature, with the x-axis showing anonymized coding of patient samples ([57], see Suppl. Information). The three patient groups (HC, IFNneg SLE, pos SLE) are color coded as in Fig 1A. **B,** Heatmap of RNA-seq data with PMN samples, but showing differentially expressed TE families. **C,** Dot plot of pathway analyses applying the indicated classifications to GSEA results for each patient subgroup. **D,** Number of genes, ISGs and TE’s for which differential expression achieved P_adj_ ≤ 0.05 in each indicated pairwise comparison, ISGs and TEs in each comparison. Note that most TEs and ISGs show predominantly increased expression.

The enrichment of the interferon gamma response observed in GSEA of IFNpos compared to HC or IFNneg samples suggested the involvement of T cells or NK cells, the major IFN-γ-producing immune cells. However, T cells from the IFNpos subset of SLE patients had no more *IFNG* mRNA at baseline than T cells from healthy controls, whereas those in the IFNneg subset had a modest reduction. NK cells were not examined in the prior study from which our data were drawn [57], but it would be worthwhile in future studies to compare *IFNG* mRNA and interferon-gamma protein expression in NK cells from IFNpos and IFNneg SLE patients and healthy controls.

### Relationships between TE expression and clinical disease

As a chronic disease whose activity varies across time and patient visits, and which is of different overall severity in different patients, the clinical status of SLE patients can be scored either as a “SLE disease activity index” (SLEDAI), on an integer scale at the visit when blood was collected, or binned according to severity (mild, moderate, severe). SLEDAI scores of 1-5, 6-10, and 11-20 indicate mild, moderate and high disease activity at the time of the visit, evaluated based on clinical parameters such as arthritis, vasculitis, myositis, pericarditis, hematuria and so on, whereas disease severity is a more subjective assessment of overall disease summed over multiple visits. These two metrics tended to correlate in the patient cohort used for our analysis (**Fig 3A**) [59]. To explore the relationship between TE expression and clinical SLE, we calculated Pearson’s correlation scores and performed linear regression analyses comparing the ISG gene set variance analysis (GSVA) score for each SLE patient and their SLEDAI score or overall disease severity (higher SLEDAI scores indicate more active disease at the time of the visit). When calculated for all SLE patients, there was a weak correlation of the ISG GSVA with SLEDAI (**Fig 3B**). While “severity” is not a clearly quantifiable value, individual patients’ ISG GSVA tended to increase in concert with their overall SLE severity (**Fig 3C**).

**Figure 3.**
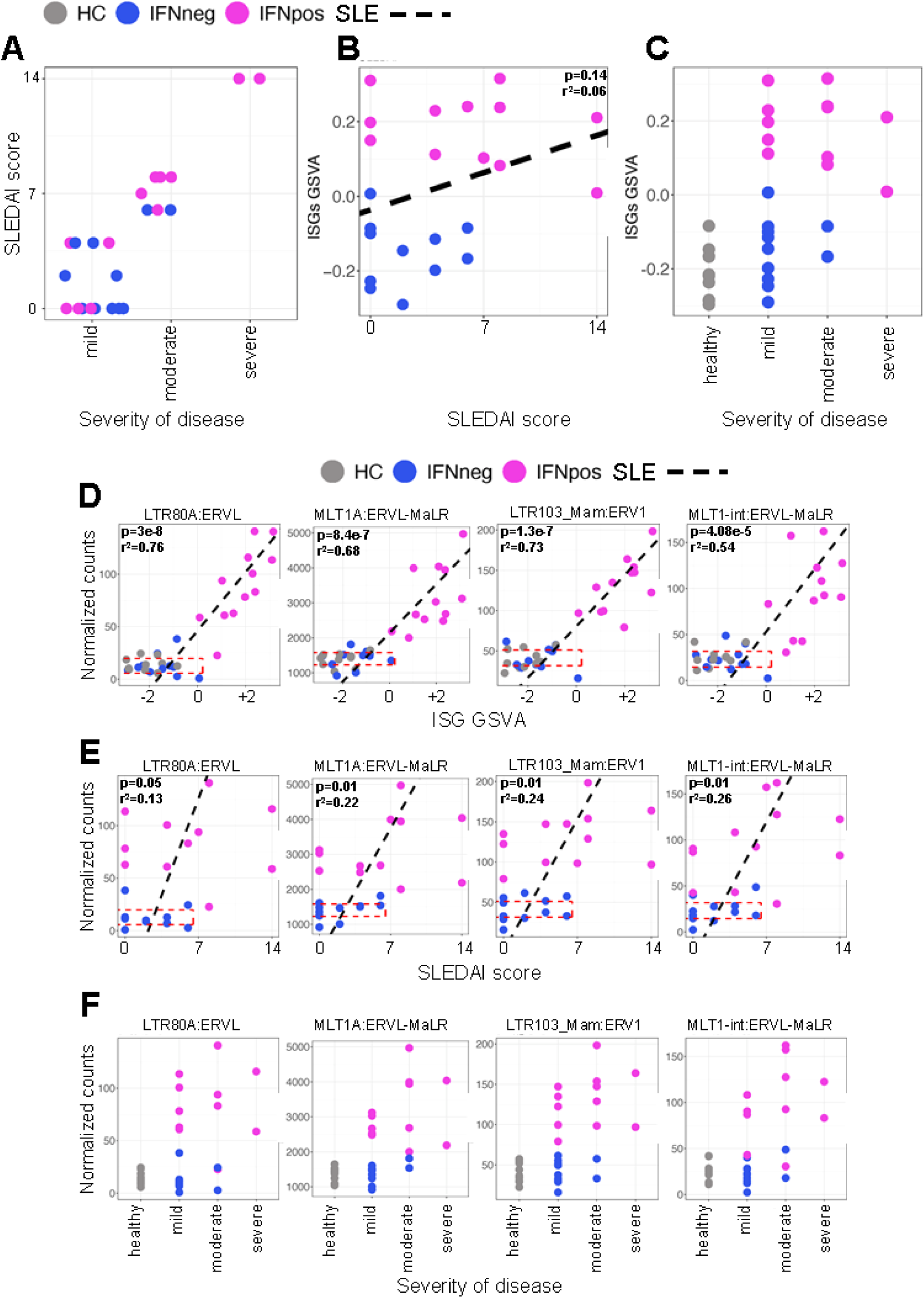
Relationship between TE increases and disease activity. A, Scatter plot of SLEDAI vs severity of disease for each of the two categories of SLE patients. **B,** For each patient sample, the ISG GSVA score in PMN is plotted in relation to the SLEDAI score, with IFNneg and IFNpos strata distinguished by color-coding. The dotted black line shows the linear regression result derived using all 23 SLE samples. **C,** Similar to B but a scatter plot of ISG GSVA scores vs severity of disease category of SLE patients and HC. **D-F,** Scatter plots focused on four specific TE families, as indicated, using data from individual patient samples. **D,** Normalized counts of the indicated TE in a subject’s PMN plotted against the ISG GSVA score in the same sample. A linear regression line calculated for all samples (black dotted line) is shown, with a rectangle delimited by a red dotted line showing the 95% confidence interval for the normalized TE counts of the HC and IFNneg SLE samples with each TE family. **E,** As in D but plots of normalized counts for the indicated TE family vs SLEDAI score. Linear regression results are denoted by dotted lines as in B. **F,** As in E except the TE counts are plotted against disease severity as tabulated in the original work [57].

We then calculated the correlation of ISG GSVA and SLEDAI scores with normalized read counts for specific TE subfamilies. This analysis identified 17 TE subfamilies that were differentially expressed in one or more of the comparisons for PMN from patient groups SLE vs HC, IFNpos vs HC, IFNpos vs IFNneg and whose expression also correlated positively and significantly with ISG GVSA and SLEDAI score (**Suppl. Fig 1C**). As examples, we show four of the top differentially expressed TE (LTR) subfamilies: (i) LTR80A:ERVL, (ii) MLT1A:ERVL-MaLR, (iii) LTR103_Mam:ERV1 and (iv) MLT1-int:ERVL-MaLR (**Fig 3D-F**). For each TE subfamily, we plotted the magnitude of TE expression (normalized read counts) for each patient’s PMN against the ISG GSVA in the same sample (**Fig 3D**). The ISG GSVA and TE counts were strongly correlated, with the IFNpos subset showing both the highest TE expression levels and (as expected) the highest ISG scores.

To explore the relationship between increased TE expression and clinical SLE, we calculated Pearson’s correlation scores comparing the SLEDAI score of each SLE patient with quantified TE expression (**Fig 3E**). For the four differentially expressed TE subfamilies listed above, TE counts correlated positively with SLEDAI score (**Fig 3E**) and severity of disease (**Fig 3F**). Thus, quantification of transcript levels for certain TE subfamilies correlates at least as well with SLE activity and clinical severity as do measures of ISG expression in the IFNpos subset of SLE patients.

Notably, the data show that the relationships noted for increased TE expression were not confined to PMN. Upregulated TE subfamilies whose expression correlated positively with the SLEDAI score could be identified in each cell type except, intriguingly, B cells (**Suppl. Fig. 3A**). Although LTR-like elements contributed the majority of severity-correlated TEs in PMN, they comprised fewer than half of such elements in classical dendritic cells (cDC) and classical monocytes (cMo) (**Suppl. Fig. 3A**). However, LTR-like TEs were the major group whose upregulation correlated with the ISG GVSA score for each cell type in the IFNpos group (**Suppl. Fig. 3B**). The differences in overall counts of TE-encoded RNA were modest, both when comparing different cell types in HC, SLE IFNneg and SLE IFNpos, and when comparing TE levels in a particular cell type between healthy controls and either IFNneg or IFNpos SLE patients (**Suppl. Fig. 3C**). In contrast, the differences in total ISG counts among different cell types were more substantial, either when analysing baseline ISG expression in different cell types of healthy controls or comparing ISG levels in a particular cell type between healthy controls and IFNpos SLE patients (**Suppl. Fig. 3D**). We conclude that although there was substantial upregulation of specific TEs in each leukocyte type analyzed in this patient group, their increased expression led to modest changes in the overall load of TEs. This in turn implies that only a small fraction of total TEs show upregulation in SLE, suggesting a stochastic process rather than a broad upregulation affecting all members of a TE family or subfamily.

We further investigated the relationship between expression of ISG genes and that of their intronic TEs, considering the variation of these expression levels across different cell types. For each intronic TE of an ISG, we computed the correlation of that TE’s expression with ISG expression across the six cell types and deemed this significant if correlation value was greater than zero with a p-value less than 0.05 (**Suppl. Fig. 3E; Suppl. Table 3**). If TE expression were mainly explained by ISG expression, one would expect most or all TEs within an ISG to have a highly correlated expression with that of ISG. Indeed, for 3.13% of ISGs (21/672), expression of half or more of all the TEs correlated with ISG expression (bars between 0.5 and 1.0); similarly, for 8/672 ISGs (1.19%), all TEs correlated with ISG expression (bar at 1.0) (**Suppl. Fig. 3E; Suppl. Table 3**). In contrast, expression of the largest fraction of ISGs (433/672, 64%) was correlated with expression of intronic TEs across all cell types (>0 and ≤25%). These results suggest that increased expression of an intronic TE cannot be explained simply by increased expression of the ISG within which the intronic TE is contained.

### Relationship of intronic TE expression to expression of surrounding genes in SLE

Quantification of the expression of uniquely mapped TEs showed enrichment of SINEs, LINES, LTRs and DNA elements in all SLE patients compared to HC, as well as in IFNpos SLE samples compared to HC and IFNneg SLE (**Suppl. Fig 1D**). We classified the genomic location of these differentially expressed TEs (DE TEs) as (a) within 2 kb upstream of a nearby gene, (b) in exons, (c) introns, (d) 3’-UTRs, (e) 5’-UTRs, and (f) intergenic regions (**Fig 4A**). Most of the DE TEs in the pool of all SLE samples (IFNneg and IFNpos) mapped to introns (blue bars) of differentially expressed genes (DEGs; cluster labelled TE UP, GENE UP), although a few DE TEs were also observed in the introns of genes that were not differentially expressed (**Fig 4B**,TE UP, GENE NO; for actual frequencies and numbers see **Suppl. Fig. 4**). This pattern was also observed for the subgroup of IFNpos SLE samples (compared to healthy controls or IFNneg samples; **Fig 4C, D**), but the stratification increased the sensitivity so that 4-to 7-fold more DE TEs could be assigned to genomic positions. Consistent with **Fig. 1H and 2D**, there was virtually no signal for DE TEs in IFNneg SLE PMN samples compared to controls.

**Figure 4.**
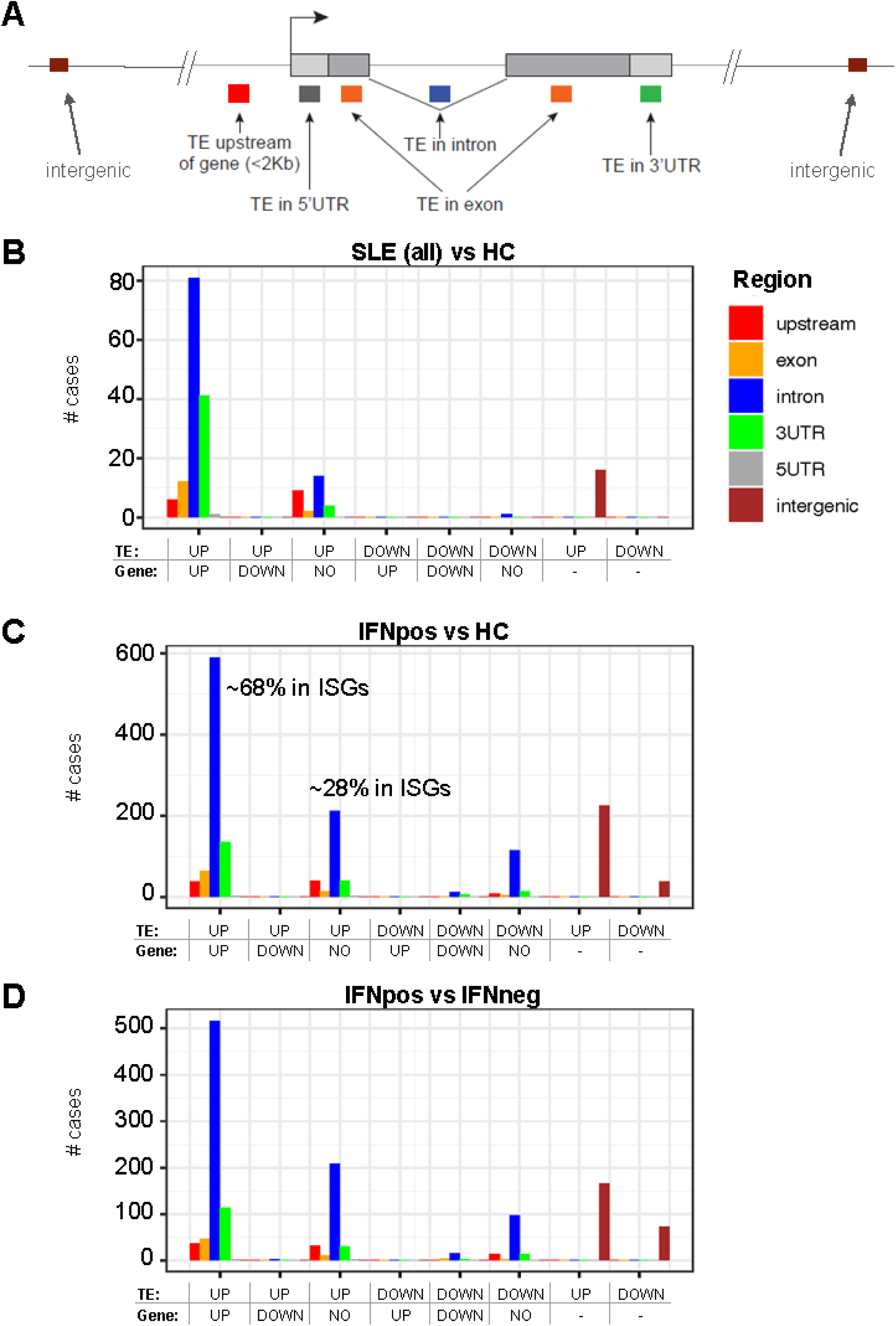
Genome annotations for locations of DE TEs relative to conventional genes. A, Representation of the potential locations of a TE relative to a gene. **B-D,** Bar graph y-axes show the number of differentially expressed (DE) TEs counted for each type of genomic location denoted in A, subdivided based on whether the RNA counts for the TE increased or decreased, and then by whether expression of the linked gene increased (“UP”), decreased (“DOWN”) or did not change signficantly (“NO”). **B,** Shown are the numbers of DE TEs mapping to each type of genomic location relative to identifiable genes when analyzing DE TEs identified by comparisons of all 23 SLE samples to HC. **C,** As for B but using the DE TEs identified by comparing HC samples to IFNpos SLE PMN. The inset percentages are the fraction of genes for which TE’s specifically mapped to an intronic region of an ISG. **D,** As for B but using the DE TEs identified by comparing RNA-seq of IFNpos and IFNneg samples.

Expression of genes associated with upregulated TEs was generally increased (∼80% of instances), and sometimes unchanged (∼20% of instances), but never decreased; in no instance was the direction of change the opposite for DE TEs and the linked gene (i.e., gene UP, TE DOWN or vice versa) (**Fig. 4B-D**). The majority of upregulated TEs were present in introns or 3’ UTRs of genes whose expression also increased (TE UP, GENE UP); ∼68% of the upregulated genes were ISGs (**Fig. 4C**; for details see **Suppl. Figs. 5A, 5B**). These data suggest a positive correlation (perhaps stemming from a mechanistic connection) between increased expression of TEs and increased expression of their associated genes, which could potentially reflect intron retention or TE exonisation. In over 200 instances, however, expression of an (intronic) TE was increased and yet that TE mapped to the vicinity of a gene with no change in expression (TE UP, GENE NO); only ∼28% of these unchanged genes were ISGs (**Fig 4C, D; Suppl. Fig 5A, B**). In particular, in SLE compared to control PMN, increased expression of intronic LINE1 (L1) elements was more associated with genes whose expression was unchanged (**Suppl. Fig. 5C**).

**Figure 5.**
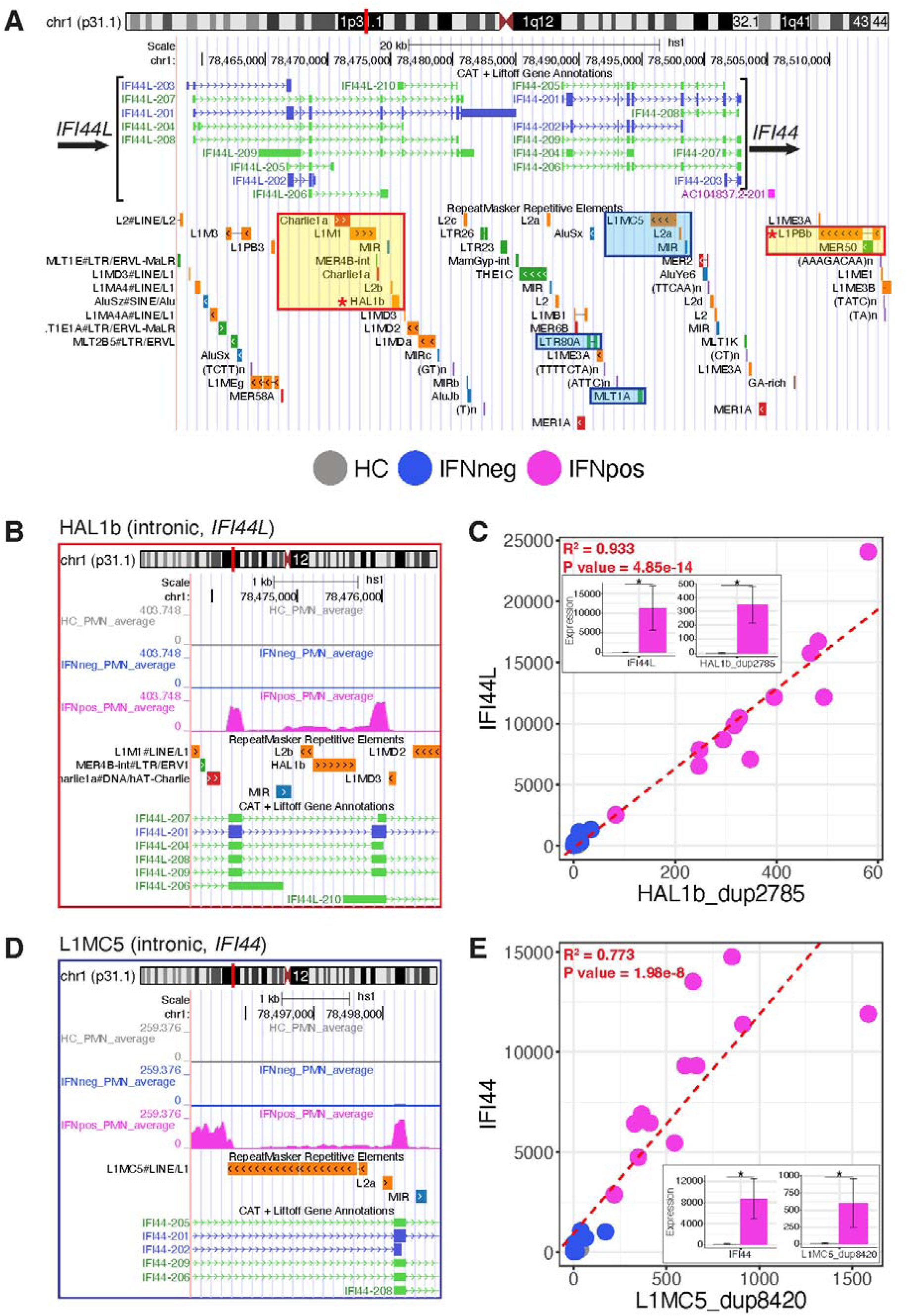
Expression of HAL1B and L1MC5 intronic LINE-1 elements in the *IFI44L* and *IFI44* genes correlates with increased *IFI44L* and *IFI44* gene expression and with splicing variations in adjacent exons. A, Genome browser view of the *IFI44L-IFI44* locus. In order, the tracks depict chromosomal location, the annotated splice isoforms of the *IFI44L* and *IFI44* genes, and all repeat/transposable elements in the *IFI44L-IFI44* locus. The HAL1b (panel B) and L1MC5 (panel D) units whose expression is increased in IFNpos SLE PMN are denoted by orange or blue boxes. The intergenic L1PBb_dup129 (3’ of *IFI44*) and the intronic HAL1B_dup2785 elements (indicated by asterisks in the boxed enclosures) were the most highly upregulated individual elements within their respective subfamilies. **B-D**, zoomed-in genome browser views showing averaged RNA-seq reads for all samples in each of the patient groups – healthy controls (HC; gray), IFNneg (blue), and IFNpos (mauve). For **B** and **D**, note the correlation of intronic transcription encompassing the TE with splicing variations in the flanking exons. **E**, Bar graph showing averaged RNA expression for the indicated genes and their corresponding intronic TEs (*top*, *IFI44* and L1MC5_dup129; *bottom*, *IFI44L* and HAL1B_dup2785). Normalized read counts from nonstranded RNA-seq +/-standard deviation are shown. For additional information please see **Suppl. Figure 7**.

The connection of upregulated TEs to introns of upregulated genes was not limited to ISGs. Rather, the numbers and expression levels of intronic TEs were similar for ISG and non-ISG genes and gene sets (**Suppl. Fig 5D, E**); ISG genes and non-ISG genes had similar intron length distribution. Further, for ∼25% to ∼50% of upregulated TEs that mapped to introns containing multiple TEs, less than a quarter of the TEs within the intron were increased in expression (**Suppl. Fig. 5F**). These findings make the case that a meaningful fraction of DE TEs are regulated autonomously from the surrounding gene in PMN of SLE patients. While more extensive long-reads sequencing studies are needed to establish this point unambiguously, the detection sensitivity for upregulated TEs associated with genes whose expression is unchanged (TE UP, gene NO) can be increased up to 10-fold by focusing on data from IFNpos SLE subjects (**Fig 4B** *vs* **Fig 4C, D**).

Our finding that upregulated TEs were commonly located in introns of expressed genes (**Fig. 4; Suppl. Fig. 5A**) raised a general question about the molecular basis for increases in intron-derived TE-encoded RNAs in SLE. In principle, increased expression of intronic TEs might reflect transcription through the intron in which the TE was located (’bona fide’ intron retention (IR) due to regulated alternative splicing) or co-option of a TE as an alternative exon of an expressed gene. Indeed, intron retention was proposed to be increased in analyses of circulating leukocytes of SLE patients [63]. More trivially, however, apparent TE upregulation can be due to batch effects and stem from different levels of contaminating intron-containing or unspliced nuclear RNA in different samples [64]. Quantification of intron retention in the analyzed samples using the IRFinder software (**Suppl. Fig. 6**) indicated that there were very few instances in which there was a substantial likelihood of intron retention (**Suppl. Fig. 6E, G, H**), suggesting that global intron retention or batch effects were not major mechanisms accounting for increased TE expression in SLE PMN.

Nonetheless, detailed inspection of individual TEs suggested that high intronic TE expression correlated in many cases with upregulated expression of interferon-induced genes, as well as with splicing alterations in nearby exons of the gene (**Figure 5; Suppl. Fig. 7-9**). To illustrate, several upregulated elements – HAL1B_dup2785 located in an intron of *IFI44L* (**Fig. 5A**, *left yellow rectangle with red border, highlighted with asterisk*) and L1PBb_dup129 located 3’ of *IFI44* (**Fig. 5A**, *asterisk, right yellow rectangle*), and the L1MC5 and LTR80A elements mapping to introns 5 and 2 of *IFI44*, respectively (**Fig. 5A**, highlighted with blue rectangles) – were contained in regions with high TE density in the *IFI44L-IFI44* locus. Other examples in the *IFI44*, *IFIH1* (encoding MDA5), *IFI6* and IFI44L loci are shown in **Suppl. Figs. 7-9**. In each case expression of the intronic TEs (i) was of considerably higher magnitude in IFNpos compared to IFNneg or healthy control PMNs; (ii) was observed in every IFNpos sample, albeit to different degrees; (iii) correlated strongly with increased expression of the ISG that contained the expressed TE; (iv) was lower in magnitude than expression of the exons of the ISG in which the TE was contained (**Fig 5B-E; Suppl Figs 7, 8**). Generally, only a portion of the intron containing the expressed TE was transcribed, perhaps arguing against batch effects involving contamination with unspliced nuclear RNA (note that polyA+ RNA sequencing was performed in ref [59]) or canonical intron retention which might be expected to involve transcription of the entire intron. This conclusion needs to be confirmed by long-reads sequencing, however.

Genome browser views of annotated exons in the vicinity of upregulated intronic TEs showed splicing variations in annotated exons adjacent to the upregulated TEs. The mechanism of this effect is unknown, but recalls reports of intron readthrough and TE “exonisation” in cancers and primary human macrophages [32, 42, 43]. Overall, our data on intronic TE expression in IFNpos PMN in SLE emphasizes (i) the preferential enrichment of upregulated intronic TEs in introns of ISGs versus non-interferon-stimulated genes; (ii) the strong correlation of ISG upregulation with increased TE expression; and (iii) the potential association of upregulated intronic TEs with splicing changes in ISG exons flanking those TEs. We note, however, that our identification of several hundred uniquely identified intergenic TE sequences located some distance from any annotated gene (nearly 20% of all reads in TEs; **Figs. 4C, D, Fig. 6**) argues against intron retention or batch effects as the only mechanisms for increased TE expression. Moreover, increased expression of these intergenic TEs was higher in PMNs of most IFNpos compared to IFNneg SLE patients or healthy controls (compare **Figs. 6A**, **6B**). Based on these data, we suggest that mechanisms independent from intronic retention contribute a meaningful fraction of the increased TEs in PMN of SLE patients.

**Figure 6.**
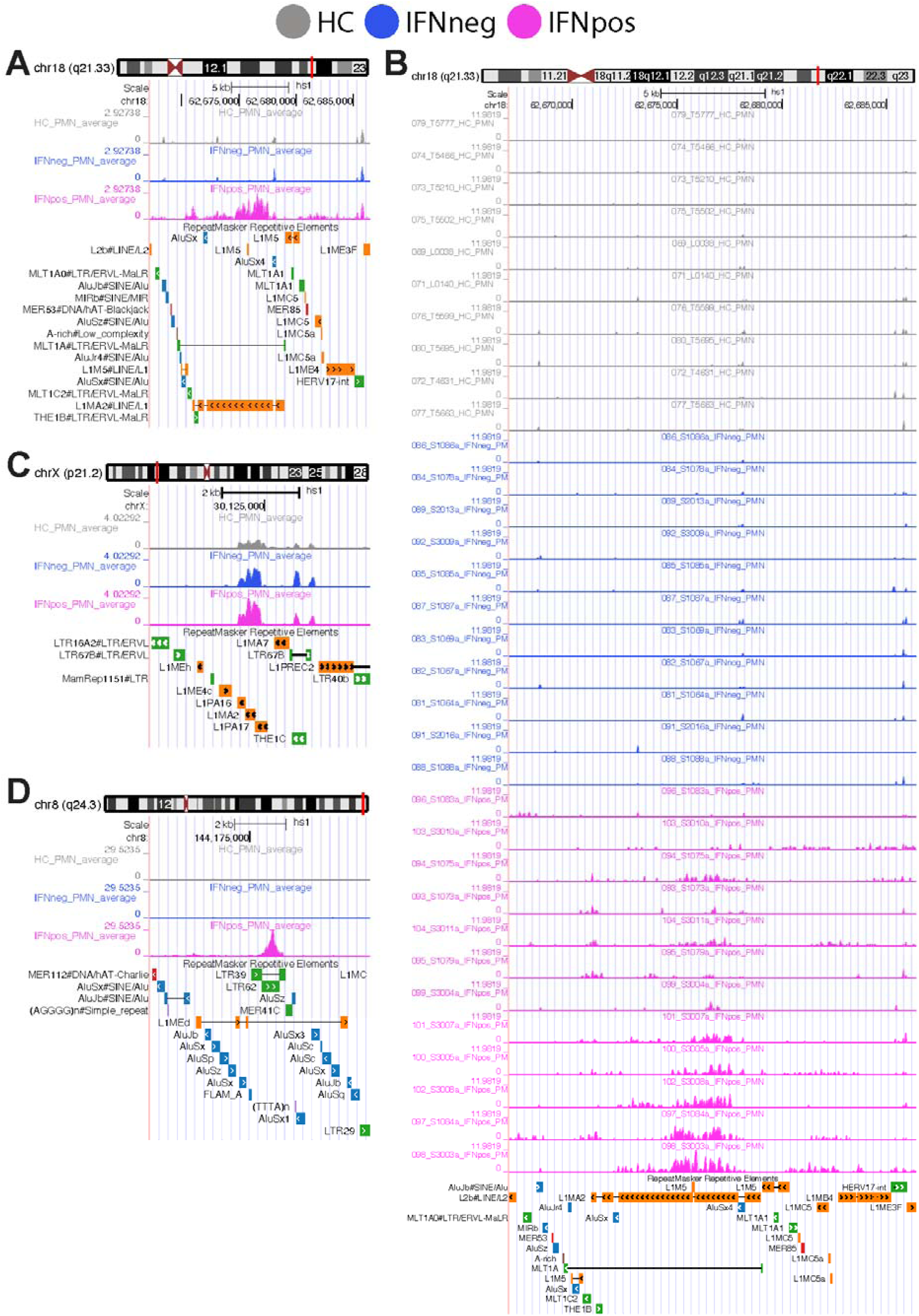
Intergenic individual copies of TEs activated as independent transcriptional units. **A**, Genome browser view of concentrated averaged signal in intergenic L1MA2 element. **B**, Expanded genome browser view of concentrated individual sample signal in intergenic L1MA2 element. **C**, Genome browser view of concentrated averaged signal in intergenic L1PA16, L1MA2 and L1PA17 element. **D**, Genome browser view of concentrated averaged signal in intergenic LTR62 element.

## DISCUSSION

Although this has not been observed in every case [20], increased expression of TE-encoded RNA can often be observed in circulating leukocytes from non-stratified collections of SLE patients [51–58, 66, 67]. Differential TE expression has been noted in CD4^+^ T cells, CD14^+^ monocytes, CD19^+^ B cells, and natural killer cells from peripheral blood mononuclear cells of SLE patients [66]. Most TE classes reported as upregulated have been in the LTR class (e.g. [19]), but a subset of PMN in SLE patients express Orf1p and Orf2p proteins encoded by L1 retrotransposons [54]. We show here that within multiple leukocyte types of a sample population of SLE patients, expression of TE families above the levels measured in samples from healthy controls was restricted to patients pre-classified as IFNpos based on the previously reported gene network analysis [59]. In contrast, none of the samples from IFNneg patients displayed such increases (**Figs. 1, 2; Suppl. Fig. 2B**). The expression levels of several differentially expressed subfamilies of highly expressed TEs in the patient population tracked closely with ISG expression and with the clinical activity and disease severity of SLE (**Fig. 3; Suppl. Fig 1C, Suppl. Fig. 3A, B; Suppl. Fig 9**). Individually-mapped, highly-expressed TEs tended to reside in the introns of genes (particularly ISGs) that were also highly upregulated in the IFNpos SLE population (**Fig. 4**). Of note, however, this correlation was not restricted to ISGs; in contrast, TE increases were also observed in upregulated exon-intron genes outside the ISG class **(Suppl. Fig. 5D, E**). The expression of some (but not all) of these TEs correlated with readthrough transcription (partial intron retention) of the corresponding introns, as well with the presence of annotated splice variations in the adjacent (flanking) exons (**Fig. 5; Suppl. Fig. 7-9**). Moreover, the increased sensitivity afforded by stratification between IFNpos vs IFNneg samples, and a focus on PMN as the cell type with highest TE upregulation, allowed us to determine that substantial fractions of the populations of upregulated TEs mapped either to genes whose expression was not increased, or to intergenic regions (**Fig. 4C**). Taken together, the findings provide evidence in support of models in which multiple independent mechanisms, and distinct categories of TE localisation (5’ UTR, exon, intron, 3’UTR, >2kb distant) relative to annotated exons and introns of genes, combine to yield the landscape of altered TEs in WBC of SLE patients.

After identifying the prevalence of TE dysregulation in each evaluable leukocyte type, neutrophils (PMN) were selected as a focus for this work in part because the overall number of differentially expressed genes, and the number of those with ISG identities, were largest in this cell subset in the data set [59] on which the analyses were based (**Fig. 1B**). Abnormalities of neutrophils are widespread among SLE patients (reviewed in [8]), and neutrophil perturbations in SLE include heightened expression of multiple cytokines, including BAFF and type 1 IFN, that can promote the breaches of self-tolerance among B cells that lead to auto-Ab production [18, 60, 61, 67]. Circulating granulocytes of SLE patients include the low-density granulocyte, a variant of typical PMN [7, 8, 54, 69], suggesting that it would be worthwhile in future investigations to distinguish this PMN subset from conventional PMN. PMN also have a specific capacity to undergo ’NETosis’, a form of programmed death that can be enhanced by PMN exposure to type 1 or 2 IFNs [70, 71]. An additional stimulus to NETosis that involves TE derepression may involve auto-antibodies directed against TE-encoded proteins. Immune complexes formed with circulating antibodies directed against antigens encoded by ERV-K102 have been reported to stimulate NET formation in vitro [58], with the caveat that immune complexes with other types of antigens may also induce NETosis [69]; the physiological concentrations of these ERV-K102-containing complexes are likely to be low and their impact at these presumed low concentrations remains to be established. Antibodies directed against the LINE L1-encoded ORF1p protein are common in SLE patients, but their presence does not correlate with disease activity in all patient cohorts [67]. Whatever the mechanisms of sensitization and triggers of NETosis, the ribonucleoproteins of neutrophil extracellular traps (NETs) can stimulate IFN-I production [7, 73], and the cytoplasmic and nuclear remnants of PMN NETosis provide self-antigens and stimulate the generation of auto-antibody-producing B cells that perpetuate a destructive cycle [9, 10, 69].

Our analyses of selected TE families in individual patient samples indicated that increased TE expression could be positively correlated with the presence of an IFN signature as well as with disease activity and severity in PMNs. For certain TE subfamilies, the correlation of increased TE expression with SLEDAI score and disease severity was even stronger than the correlation between SLEDAI score/ disease severity and increased ISG expression (**Fig. 3**). Since neutrophils constitute nearly half of the blood cell population, the relation of SLE disease severity to TE expression, neutrophil activation and IFN production in PMNs warrants further investigation. When ranked at the family level (**Fig. 3**), the most common differentially expressed (primarily upregulated) TEs were endogenous retrovirus (LTR/ERV)-related sequences, a finding generally consistent with prior reports in which analyses were performed with diverse mixtures of cell types [20; 58-60] or that did not examine PMNs [63]. However, our strategy of stratification to IFNpos patients, our focus on a single immune cell type (PMNs) and the use of the Telescope informatics tool [74] allowed us to identify differentially expressed RNA-seq reads assigned to all the varied classes of TE – HERVs, DNA transposons, LINEs, SINEs, and satellite repeats. Similar analyses are in process for other immune cell types.

Distinguishing the mechanisms that represent individual inciting events versus the diverse cellular and molecular processes that perpetuate disease is an intriguing issue in SLE research. Our data underscore the complexities in that upregulation or de-repression is observed for multiple families of TE. Consistent with the overall state of the literature on TEs in human SLE, the evidence is correlative and identifies the nature of altered regulation of specific elements, but direct experimental evidence supporting causality is absent. Previous studies have suggested that increased TE expression contributes to SLE pathogenesis [57, 58], and GWAS studies have implicated ISGs and many other immune-related genes in SLE [50, 51] but whether TE expression is a root cause or an effect (or both) of the abnormally heightened ISG expression observed in most SLE patients remains unclear. On the one hand, there is compelling evidence that TE expression can evoke type I IFN responses that result in ISG expression; specifically, double-stranded RNAs generated by transcription of inverted Alu repeats activate the dsRNA sensor MDA5, especially in the absence of the RNA editing enzyme ADAR1 or the presence of gain-of-function MDA5 variants associated with autoimmune disease [47, 48, 75, 76]. MDA5 forms long filaments on long stem loops of double-stranded RNA, resulting in downstream activation of the IRF3/7 and NFκB transcription factors which initiate type I interferon production and transcription of ISGs [75]. MDA5 is encoded by the *IFIH1* gene, which is one of 10 SLE “hub” genes (*IFI35, IFIT3, ISG15, OAS1, MX2, OASL, IFI6, IFITM1, IFIH1, IRF7*) identified by analysis of microarray data [65]; 9 of these (including *IFIH1*), are ISGs and the tenth, *IRF7,* encodes the interferon regulatory factor IRF7 that elicits the type I IFN response [65, 75]. GWAS studies have also implicated the Toll-like receptor TLR7 in SLE [50, 51]; TLR7 trafficked into endosomes recognizes viral and other single-stranded RNAs [77]. However, whether or not these mechanisms are triggered in patients by the selective increases in some TEs is not clear. Of note, we observed only modest changes observed here in the total counts of TE-encoded RNA (**Suppl. Fig. 3D**). Nonetheless, it remains possible that the load of certain specific dsRNA structures increases far more that what would be estimated by total RNA. Alternatively, although full-length L1 retro-elements that are capable of transposition are a very small subset of all such TEs [29, 34, 78, 79] it is possible that at least in some patients, dysregulation of the expression of L1-encoded ORF2p reverse transcriptase might mediate an LTR-based increase in immunostimulatory RNA or RNA:DNA hybrids [53, 54, 79].

Whether ISG expression promotes increased TE expression was even less clear. We show here that highly expressed TEs are enriched in the introns of genes that are differentially upregulated in SLE; this is particularly apparent when comparing IFNpos SLE patients versus healthy controls (GENE-UP TE-UP category **in Fig. 4**, **Suppl Figs. 3, 4**). Additionally, more than two-thirds of the introns with highly expressed TEs are located in ISGs (**Fig. 4**) and show moderate read-through transcription (which could represent bona fide intron retention) compared to other introns in the same ISG (**Figs. 5-7**). Notably, inspection of annotated splice variants in a limited number of cases suggested that the exons flanking ISG introns with increased TE expression and transcription were particularly prone to alternative splicing (**Figs. 5-7**). This phenomenon may be related to the widespread TE exonization reported to occur in both normal and cancer cells [32, 42, 43]. In primary human macrophages, TE exonization was shown to have functional consequences for the innate immune response – incorporation of an Alu-derived exon generated a short isoform of the type I IFN receptor IFNAR, which suppressed ISG induction by acting as a decoy receptor for type I IFNs [43]. Long-reads whole transcriptomic sequencing will be necessary to begin probing the mechanisms underlying the relation of increased ISG expression to increased expression of intronic TEs; this effort may be complicated by the fact that alternatively spliced transcripts containing exonised TEs are often [42] – though not always [43] – expressed at much lower levels (∼1%) than the canonical transcript. Follow-up studies incorporating long-read sequencing and using larger patient cohorts, and examining pre-morbid phases and progenitor cells of lupus-prone mice [80], will be required to probe early and late stages of SLE.

Dysregulation or increased expression of TEs has been observed in multiple diseases or pathological processes, particularly aging, neurodegeneration, and cancer [34, 37, 39, 81–84]. The findings raise intriguing questions about the generality of TE de-repression in inflammatory or autoimmune diseases. A number of autoimmune rheumatic diseases are distinct from SLE, but display evidence of increased type 1 or type 2 IFN levels, ISGs, and in limited literature, increased TE expression [20, 85–87]. For example, increased expression of ERV K102 and LINE-1 elements have been reported in rheumatoid arthritis as in SLE [88–91]. Both type 1 and 2 IFNs are important early in the pathogenesis of type 1 diabetes mellitus (T1D) [92, 93]. We note that most of these studies are preliminary, lacking essential cell type information and stratification (e.g. IFNpos versus IFNneg). The pace of advances in sequencing – especially the capacity for long-read sequence determination – and the development of new analytic tools make it uniquely timely to investigate such questions in depth, both in SLE and across auto-immune diseases in general.

### Limitations

Limitations of these analyses include the sample size and the pressing need for high-coverage long-read sequencing to permit more precise determination of the structure and location of intronic TE sequences and their potential role in splicing. We do not yet know the threshold levels of TE-encoded RNA, the specific structure(s) of these RNA in patients’ cells, the magnitude of increase in TE-derived RNA ’load’ needed to trigger increased type I IFN production in specific cell types of patients with different predisposing genetic factors, and the particular innate immune nucleic acid sensors involved. In addition, we do not yet know the extent to which increased TE expression is cell type-specific and whether particular subfamilies of TE preferentially trigger innate sensors in patients, whether in SLE, other autoimmune diseases, senescence, aging or cancer.

## METHODS

### Dataset

We collected publicly available bulk RNA-seq data from Panwar, B. et al. [59]. The overall dataset contains 288 samples from 91 subjects, 65 patients with SLE and 26 healthy controls (HC), with purified polymorphonuclear neutrophils (PMN) from 33 total subjects. Samples in the original study comprised six different cell types-classical monocytes (cMo) (n = 119), conventional dendritic cells (cDC) (n = 32), plasmacytoid dendritic cells (pDC) (n = 33), T cells (n = 36), B cells (n = 32), and polymorphonuclear neutrophils (PMNs) (n = 36, of which three were samples from follow-up visits). In some instances, specific leukocyte types were purified from blood of SLE patients provided by Sanguine Biosciences, Inc, as detailed in freely available Supplemental Data [59]. Based on a network analysis of the expression of interferon genes and interferon stimulated genes (ISGs), the original article identified an approach that divided the SLE patients into two subgroups, one exhibiting greater interferon effect, termed ’IFNpos’ [59] and a second subset designated as ’IFNneg’. In the present study, we adhered to this classification of the samples and terminology. Sample sources and patient characteristics (age, sex, IFN stratum) are briefly summarized in Supplemental Fig. 1A.

### RNA-seq and data analyses

TrimGalore v0.6.7 [94] was used to remove adapters, reads were mapped against the T2T-CHM13v2.0 genome reference using STAR [95] v2.7.10a (--outFilterMultimapNmax 1--alignIntronMin 20 --twopassMode Basic --alignSJoverhangMin 8 --alignSJDBoverhangMin 1 -- outFilterType BySJout --outSJfilterReads Unique --outFilterMismatchNoverReadLmax 0.04) to analyze gene expression. HT-Seq [96] v2.0.4 was used to quantify gene expression with default parameters. To analyze transposable element expression, reads were mapped against the T2T-CHM13v2.0 genome reference using STAR v2.7.10a (--outFilterMultimapNmax 100 --winAnchorMultimapNmax 100) allowing multi-mappers. Using the alignments with multi-mappers as input, we employed two programs to quantify the expression of transposable elements, TEtranscripts [97] v2.2.1 (--mode multi) was used to quantify the expression of the transposable elements at subfamily level and Telescope [98] v1.0.3 was used to quantify the expression of individual transposable elements (locus specific). Four comparisons to observe transcriptional changes between SLE samples and HC were performed: (1) all SLE sample vs HC; (2) IFNpos vs HC; (3) IFNneg vs HC; (4) IFNpos vs IFNneg using samples taken on the first visit of each patient as described in the original article. Differential expression analysis was performed in R [99] v4.2.1 using DESeq2 [100] v1.38.3. Genes with fewer than 10 read counts per group were filtered out before performing the analysis. Differentially expressed genes were defined using adjusted p value ≤ 0.05 and |log2 Fold Change| ≥ 1. In the case of TEs (at the individual and family levels), those with fewer than 1 read count in all samples were filtered out before performing the analysis. Differentially expressed TEs were defined using adjusted p value ≤ 0.05 and |log2 Fold Change| ≥ 0.5.

To analyze the relationship between the expression of transposable elements and genes, we used genomic coordinates of genes and transposable elements and used BEDtools [101] and R to identify the genomic locations of transposable elements relative to nearby genes. We classified the genomic location of the transposable elements as (a) 2 kb upstream of a nearby gene, (b) in exons, (c) introns, (d) 3’- UTRs, (e) 5’-UTRs, and (f) intergenic regions. Once we located the transposable elements relative to genes in the genome, we looked for the correlation between differentially expressed transposable elements and differentially expressed genes (Figure 3). IRFinder [102] v1.3.1 was used to quantify intron retention in the analyzed samples using T2T-CHM13v2.0 genome reference. The average expression of ISGs and the TEs within introns of those genes was calculated using the expression values of IFNpos patients for all cell types in the dataset. Briefly, for each ISG, we examined average ISG expression and average expression of intronic TEs within this ISG across all IFNpos patients for each cell type. Pairwise correlations were calculated for each gene and the intronic TEs within it using the average expression values of the genes and TEs of all six cell types. To correlate avg ISG expression across the six cell types to the intronic TE expression across the same six cell types, we computed a correlation of two vectors of size six for each ISG – intronic TE pair, taking into account that there could be multiple intronic TEs within each specific ISG.

To identify the individual copies of TEs activated as independent transcriptional units, we focused on the Differentially expressed TEs in IFNpos samples as compared to healthy controls (HC). Analyses used BEDtools [101] with the option -o mean, to determine the signal in the RNA-seq data for each TE unit and compare against the +/-1 kb upstream and downstream regions around them (noise). TEs showing at least 2-fold change in the metric “TE/noise” were manually examined using the UCSC genome browser [103].

### Gene set enrichment and variation analyses

Gene set enrichment analysis (GSEA) was performed in R v4.2.1 using clusterProfiler [104] v4.6.2 with the hallmark gene sets from msigdbr [105] v7.5.1 with the results of the differential expression analysis as input. Pathways with p value ≤ 0.05 were considered. Gene set variation analysis (GSVA) was performed in R v4.2.1 using the gsva function from GSVA [106] v1.52.3 and the interferon stimulated genes list (Supplemental Table 1) used in this study, using the list of IFNalpha or IFNgamma genes [107, 108] (Supplemental Table 2) to quantify enrichment of ISGs, IFNalpha and IFNgamma in the analyzed samples.

### Correlation analyses of transposable elements and clinical disease

Pearson correlations and linear regression were performed in R v4.2.1 using the cor.test function and the lm function, respectively, from the stats library using normalized counts of TEs from SLE patients and SLE disease activity index (SLEDAI) score to assess a possible correlation between TEs and severity of the disease or using normalized counts of TEs from SLE patients’ PMN and the ISG GSVA scores to assess a possible correlation between the expression of TEs and the expression of ISGs. TEs with positive and significant correlation were filtered based on differential expression status, those that were differentially expressed and that correlated significantly were kept for further analyses.

### Tracks

From deeptools [109] v3.5.4, multiBamSummary was used to calculate normalization factors and bamCoverage was used to generate bigWig files to explore genome coverage. Wiggletools [110] v1.2.11 and bedGraphToBigWig v4 were used to obtain average genome coverage tracks for the three conditions IFNpos, IFNneg and HC. Genome browser views were generated using the most recent update (2025) of the UCSC genome browser [103].

### SUPPLEMENTARY MATERIALS

**Supplementary Table 1. List of genes used for the ISG signature.**

**Supplementary Table 2. List of genes in the IFN-alpha and IFN-gamma signatures**. Many of these genes are included in the interferome database https://interferome.org/interferome/home.jspx

**Supplementary Table 3. Correlation analysis for expression of TEs in introns of ISGs**

## AUTHOR CONTRIBUTIONS

Data acquisition, processing via publicly available computational tools, and display in figures were performed by L.J.A.-V., H.S., and B.V.R., with input from F.A., who facilitated access to RNA sequencing and meta-data in prior published work from his lab. K.C.K. interpreted clinical data from the samples. H.S. supervised the bulk of the initial bioinformatic analyses while F.A. supervised the later stages. The project structure and evolution were conceived and directed by M.R.B. and A.R. Analyses and figure panels were designed or refined by all authors. All authors contributed to writing and editing the manuscript.

## ACKNOWLEDGEMENTS

We thank Q. Li (University of Pennsylvania) for sharing his curated list of ISGs and G. J. Faulkner (University of Queensland) for critical review of our methodology and results.

## AVAILABILITY OF DATA, MATERIALS, AND CODE

No new data or code were generated, and the means of access to freely available open sources are indicated in the Methods. No new materials were generated for this analysis. The data used in this study can be accessed in GEO under the accession number GSE149050.

## ABBREVIATIONS

SLE: systemic lupus erythematosus
Ab: antibody
IFN: interferon
type I IFN: IFN-I
ISGs: interferon signature (or stimulated) genes
TE: transposable elements
PMN: polymorphonuclear leukocytes (neutrophils)
HC: healthy controls
LTR: long terminal repeat
NET: neutrophil extracellular trap
ds: double-stranded
ERV: endogenous retrovirus
SINE: short interspersed nuclear element
LINE: long interspersed nuclear element
GWAS: genome-wide association study
GSEA: gene set enrichment analysis
GSVA: gene set variance analysis
NK: natural killer
TNFSF: tumor necrosis factor(-like) secreted factor
BAFF: B cell activating factor
DE: differentially-expressed
DE gene: DEG
UTR: untranslated region
SLEDAI: SLE disease activity index
IR: intron retention.

## HUMAN ETHICS AND CONSENT TO PARTICIPATE

Not applicable. No human subjects were used in this study, which is a reanalysis of existing publicly available datasets.

## CONSENT TO PUBLISH

Not applicable. This study is a reanalysis of existing publicly available datasets.

---

**Supplementary Figure 1.**
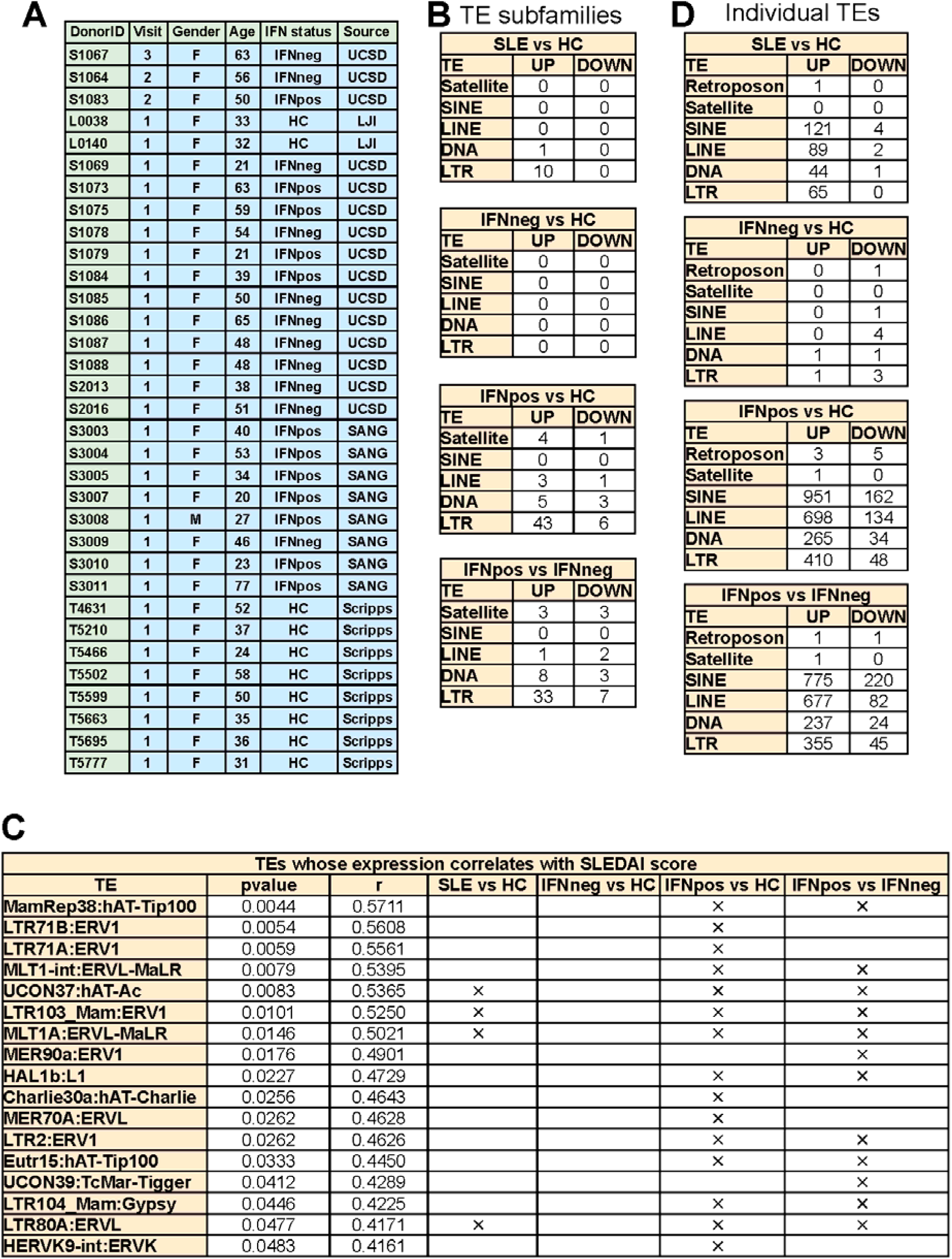
Subject characteristics, aggregate numbers of differentially expressed TE classes, and TEs with positive and significant correlation with SLEDAI score. A. Linking to the previously published Supplemental Information [59], shown are to DonorIDs of subjects whose blood samples were fractionated into the six types of leukocyte, age, sex, stratification as defined in [59], and sourcing (UCSD, Lupus Clinic of K. K.; LJI, La Jolla Institute; SANG, Sanguine Biosciences; Scripps, The Scripps Research Institute). Due to the paucity and unequal distribution of bloods from follow-up visits, all analyses presented herein were confined to samples collected at the first visit. B. Tables show the classes of TE subfamilies that are differentially expressed in PMN in each pairwise comparison between subject groups (IFNpos, IFNneg, and HC), and numbers of instances of increased expression for each subfamily. C, Table of the differentially expressed TEs that have significant positive correlation with SLEDAI score. The check mark indicates if a given TE was found upregulated in the indicated comparison. Raw p values are shown. D, A tabular enumeration of the numbers of individual differentially expressed TEs in the indicated classes and comparisons.

**Supplementary Figure 2.**
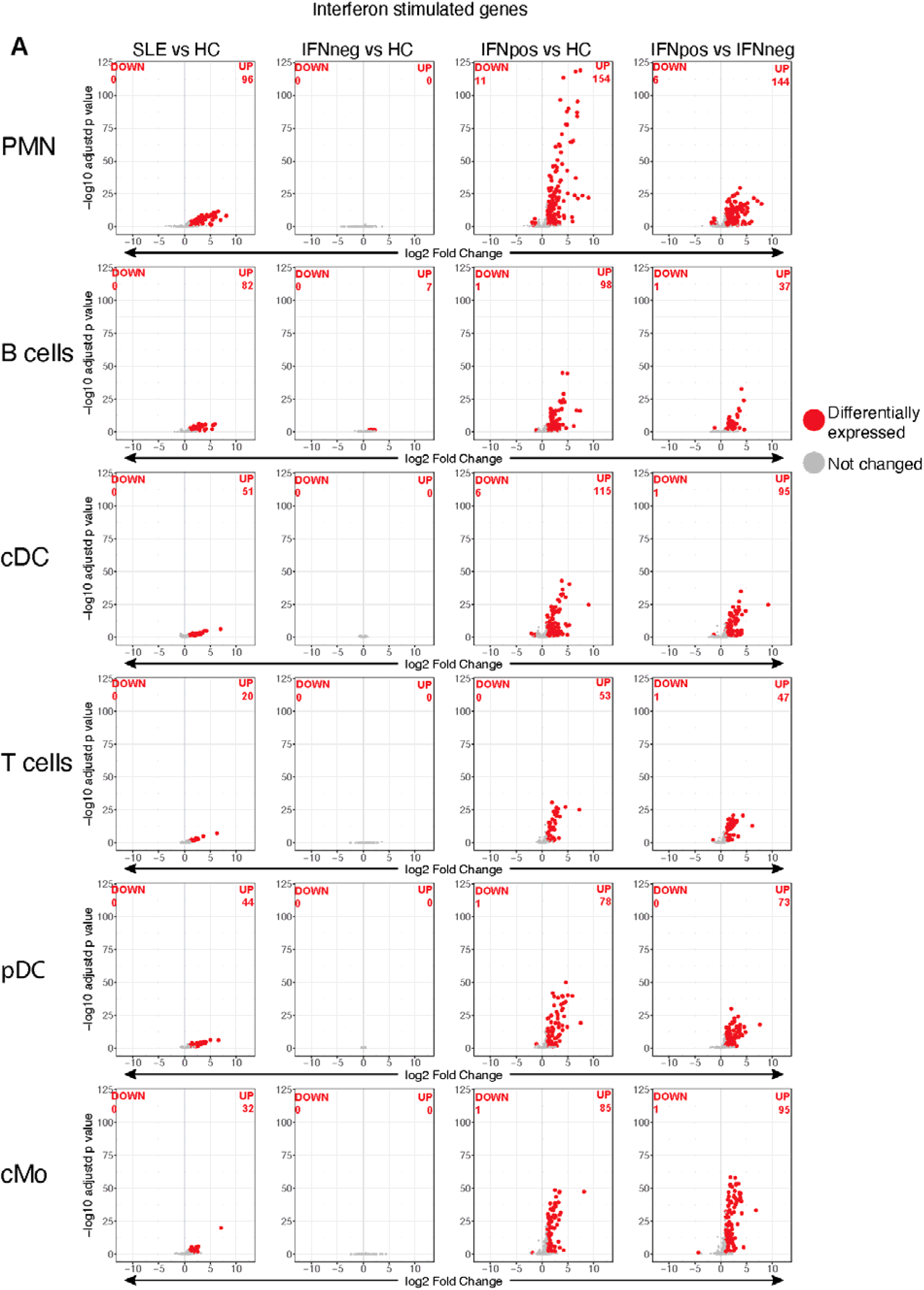
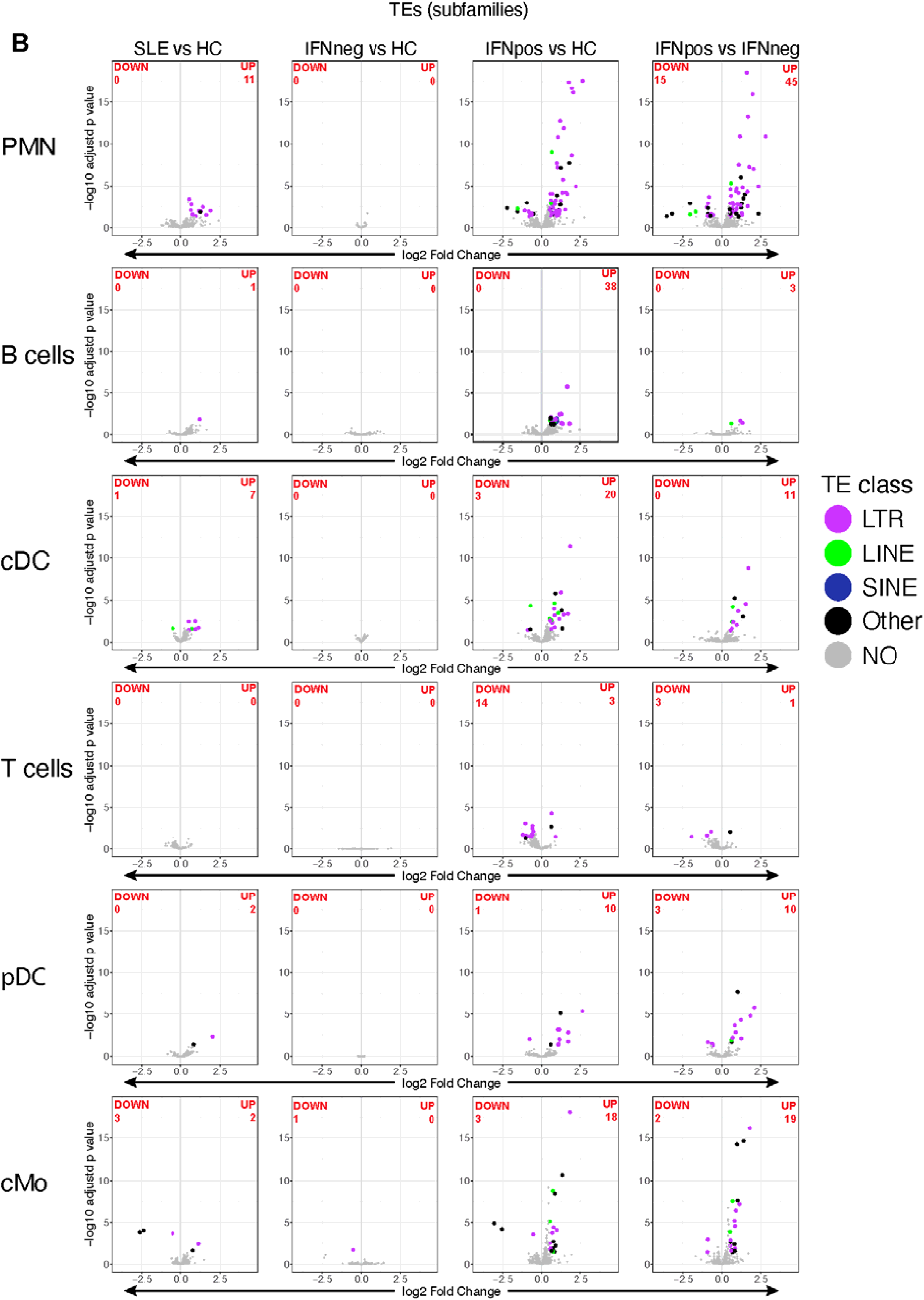
Differential expression of ISGs and TEs (subfamilies) in all cell types. For each of the six evaluable cell types (designated to the left), analyses of differential expression in pairwise comparisons between subject groups (IFNpos, IFNneg, and HC), as indicated, were performed for **A**. ISGs and **B**. TEs, classified and color-coded as indicated, using the Methods of Fig 1. Data for PMN are identical to those of Fig 1 and recapitulated to facilitate comparisons to the other types of leukocyte.

**Supplementary Figure 3.**
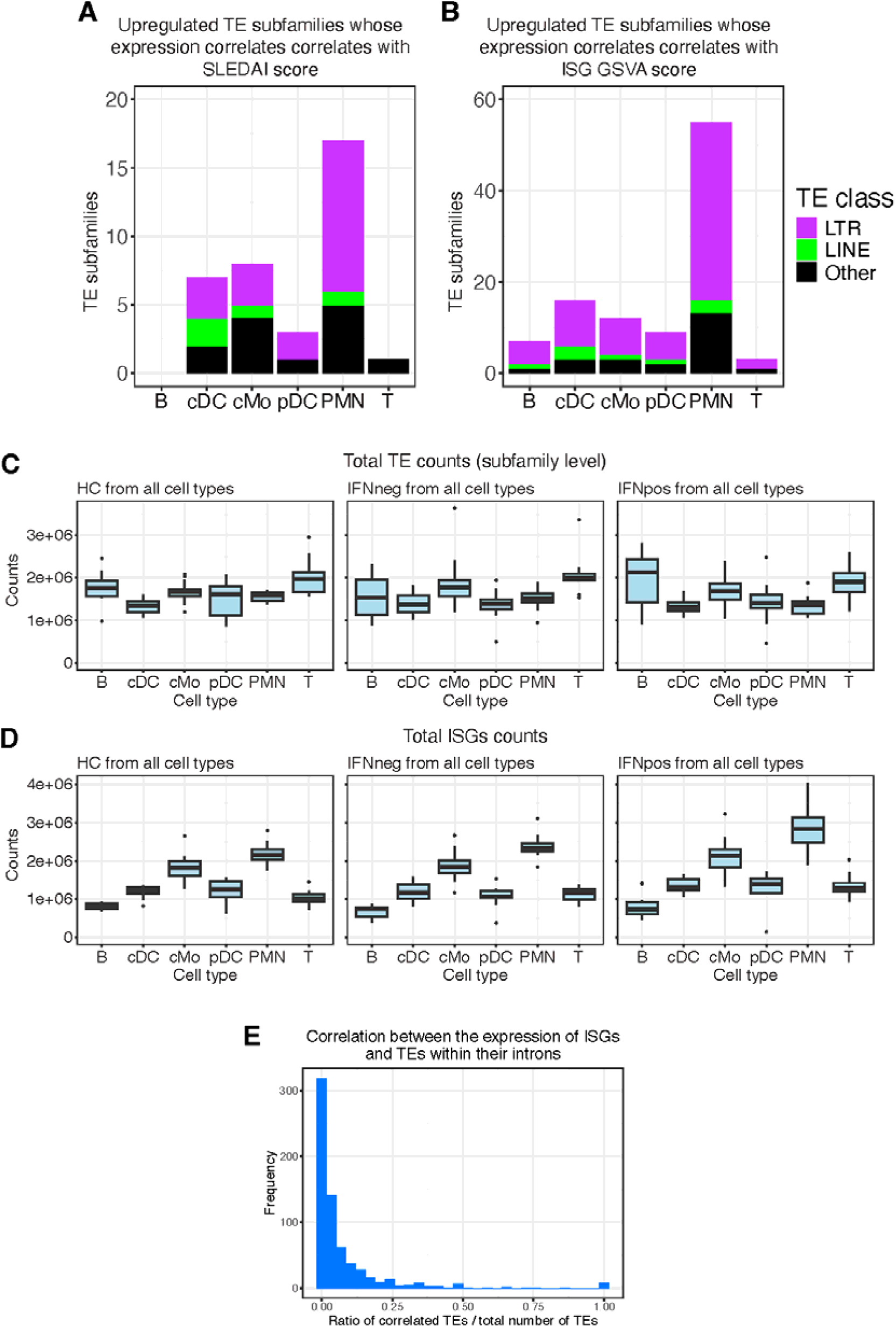
Increased expression of subsets of upregulated TEs (subfamily level) correlates with SLEDAI or ISG GSVA scores. A, For each cell type, the number of upregulated TEs (subfamily level) whose expression correlates with SLEDAI score is shown along with classification by TE class as indicated. B. As in A except the numbers are for subfamily level TEs whose increase correlated with ISG GSVA score. C, Total TE counts (subfamily level) per cell type, and D. total ISGs counts per cell type. E, Histogram showing the distribution of the ratio between the number of intronic TEs whose expression correlates with the expression of the ISG in which they are located, and the total number of intronic TEs in the ISG.

**Supplementary Figure 4.**
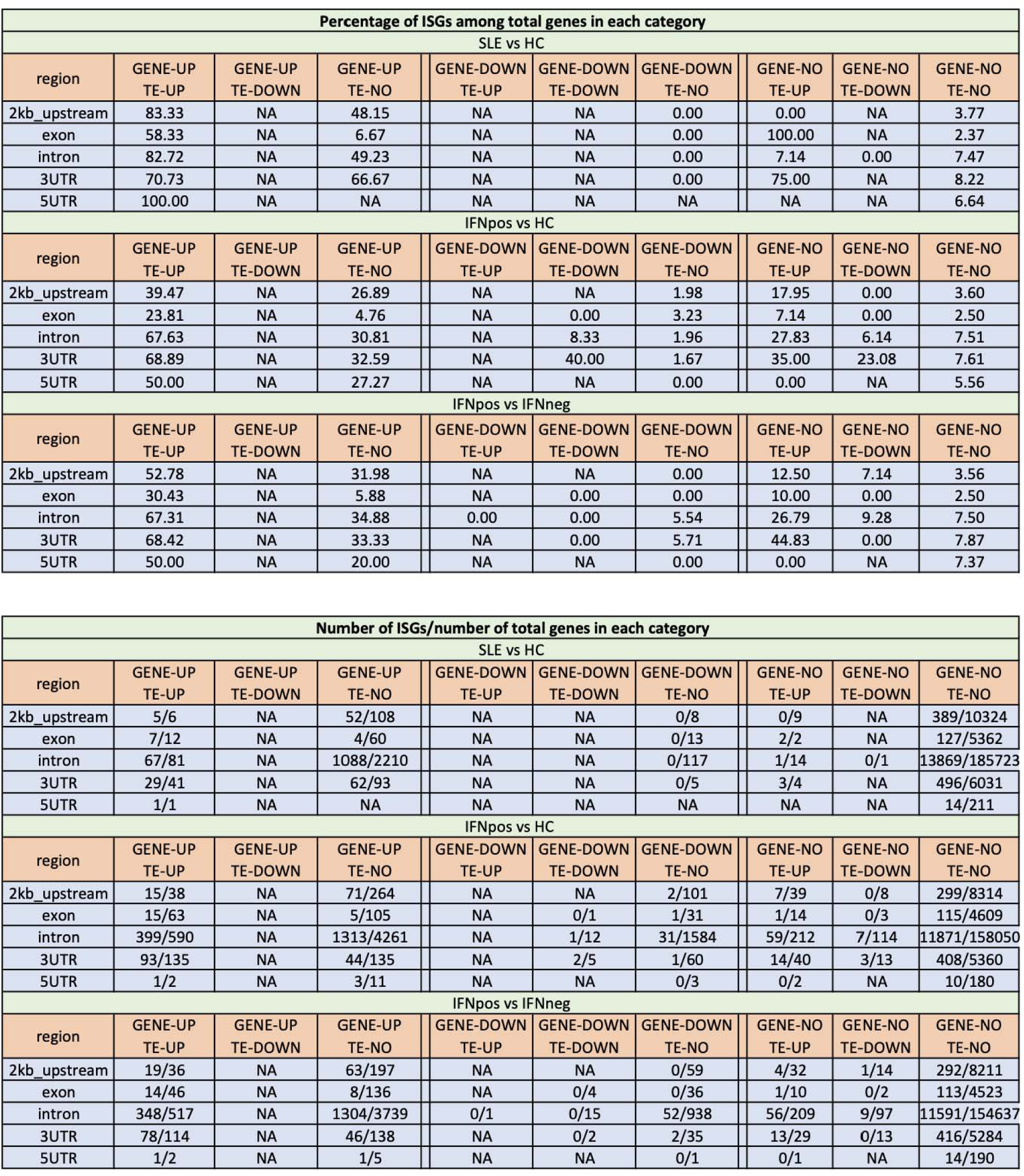
Table showing the numbers and percentages of ISGs among total genes with nearby differentially expressed (DE) TEs. Detailed quantitative breakdown of data depicted as graphs depiction in Fig. 4. For each of the indicated pairwise comparisons between PMN of two patient groups (all SLE vs HC, IFNpos vs HC, and IFNpos vs IFNneg), numerical entries in the tabulated cells indicate percentages (upper matrix) and numbers (lower matrix) of ISGs among the total of all genes containing DE TEs for the comparisons. The data are computed for TEs at each of the genomic locations shown in Fig. 4A. (N/A, not applicable).

**Supplementary Figure 5.**
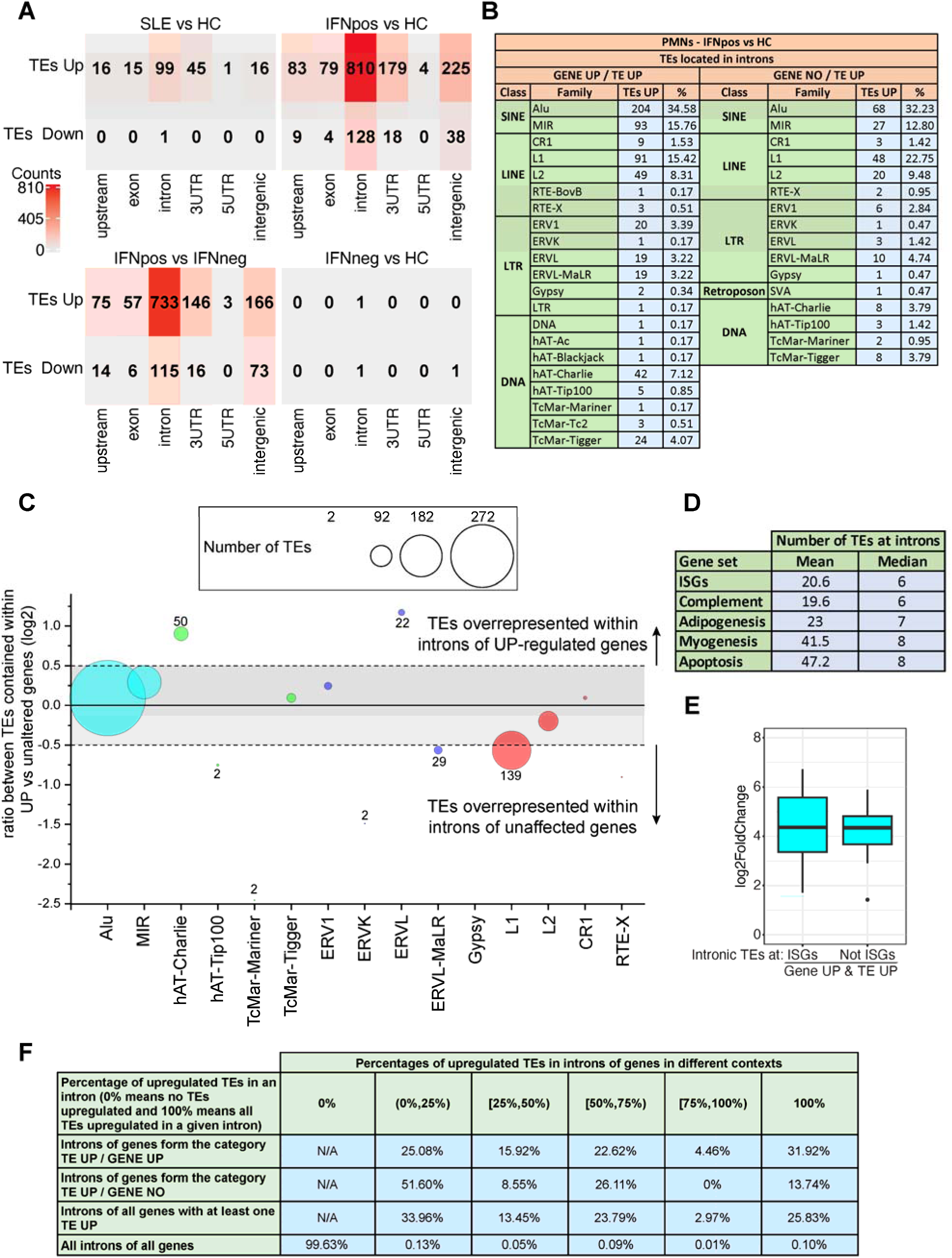
Analyses of the intronic TEs in SLE PMN relative to healthy control samples. **A**, Number of differentially expressed transposable elements found at different genomic locations (upstream, exon, intron, 3’UTR, 5’UTR, intergenic) in the comparison SLE vs HC, IFNneg vs HC, IFNpos vs HC and IFNpos vs IFNneg. **B**, Table showing the number of upregulated TEs classes and families in introns of upregulated genes (left) or not differentially expressed genes (right). **C**, Bubble plot comparing TEs families of the upregulated TEs in introns of upregulated genes and upregulated TEs in introns of not differentially expressed genes. **D**, Table showing the mean and median values of TEs in introns of different gene sets. **E**, Boxplots showing the log2 Fold Change of TEs located in introns of upregulated ISGs and not ISGs. **F**, Table showing the percentages of upregulated TEs in introns of genes in different contexts.

**Supplementary Figure 6.**
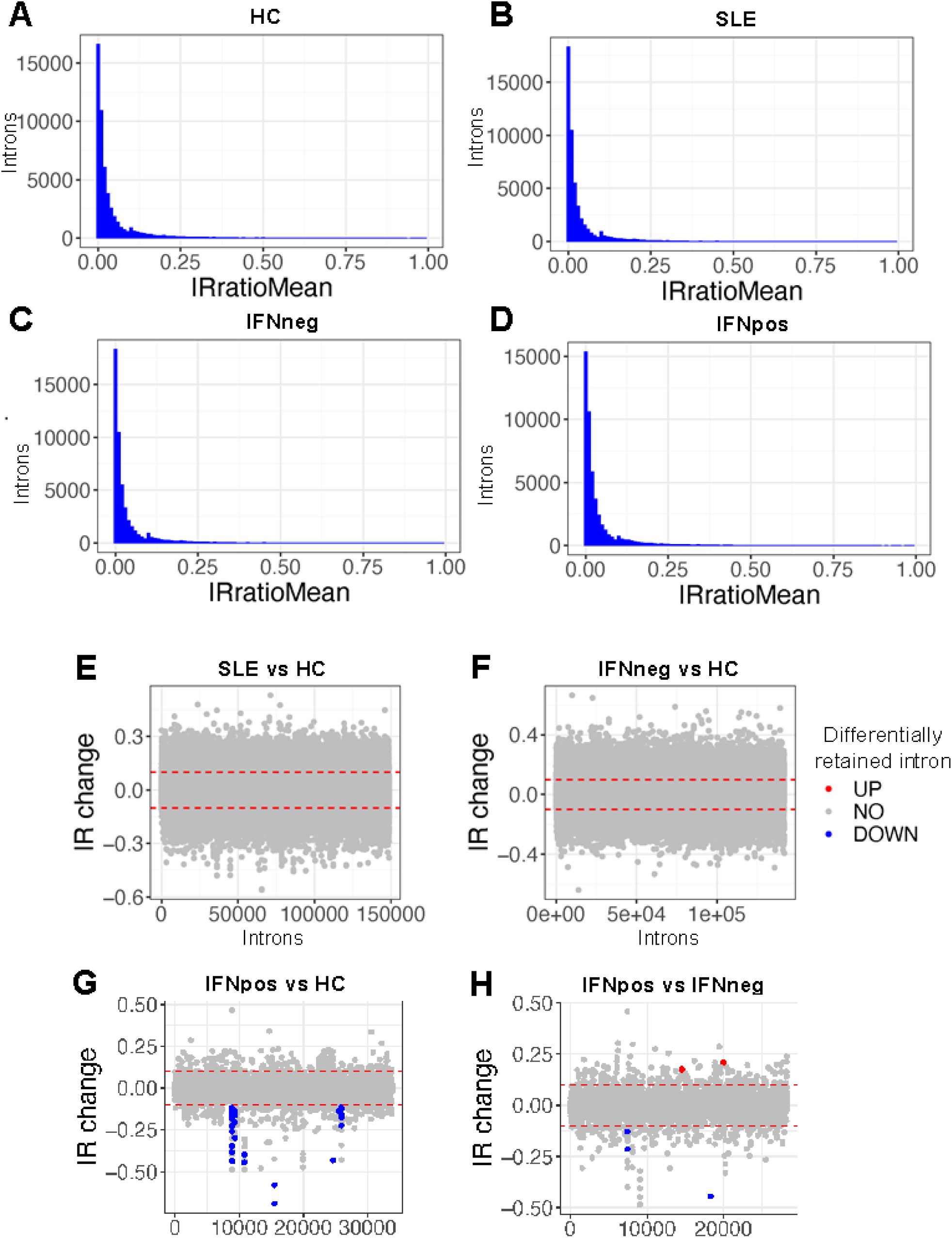
Computational estimation of the extents of intron retention in SLE PMN relative to healthy control samples. **A**, Mean of intron retention ratio (IRratioMean) from IRFinder of HC samples. **B**, Mean of intron retention ratio (IRratioMean) from IRFinder of SLE samples. **C**, Mean of intron retention ratio (IRratioMean) from IRFinder of IFNneg samples. **D**, Mean of intron retention ratio (IRratioMean) from IRFinder of IFNpos samples. **E**, Differential Intron retention comparing SLE against HC, significant intron retention change is indicated by color. Differentially retained intron in SLE (UP) red; differentially retained intron in HC (DOWN), blue; not differentially retained intron, grey. **F**, Differential Intron retention comparing IFNneg against HC, significant intron retention change is indicated by color. Differentially retained intron in IFNneg (UP) red; differentially retained intron in HC (DOWN), blue; not differentially retained intron, grey. **G**, Differential Intron retention comparing IFNpos against HC, significant intron retention change is indicated by color. Differentially retained intron in IFNpos (UP) red; differentially retained intron in HC (DOWN), blue; not differentially retained intron, grey. **H**, Differential Intron retention comparing IFNpos against IFNneg, significant intron retention change is indicated by color. Differentially retained intron in IFNpos (UP) red; differentially retained intron in IFNneg (DOWN), blue; not differentially retained intron, grey.

**Supplementary Figure 7.**
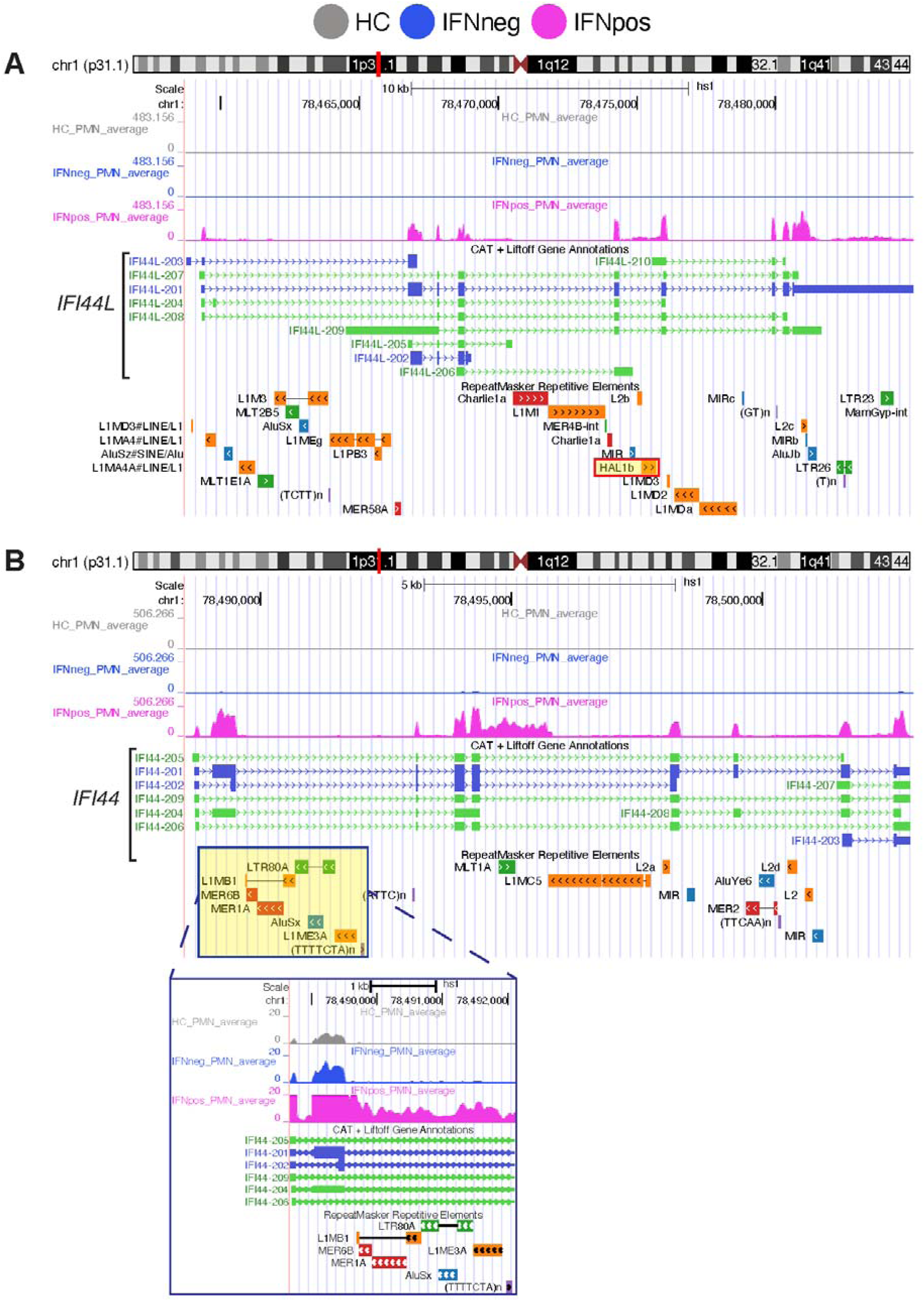
(complementary to Figure 3). Representative examples of incomplete transcription of two TE-containing introns in the *IFI44L* and *IFI44* loci. A, Expanded genome browser view of the *IFI44L-IFI44* locus showing increased expression of the same HAL1B element shown in Figure 5A, B. B, Expanded genome browser view of the *IFI44* gene, showing one of LTR80A:ERVL’s and one of MLT1A:ERVL-MaLR’s highest expressing units located in introns 2 and 5, respectively, of *IFI44*. Note the selective upregulation of transcription in the introns containing these high-expressed TEs and the occurrence of annotated splice variants in the flanking exons.

**Supplementary Figure 8.**
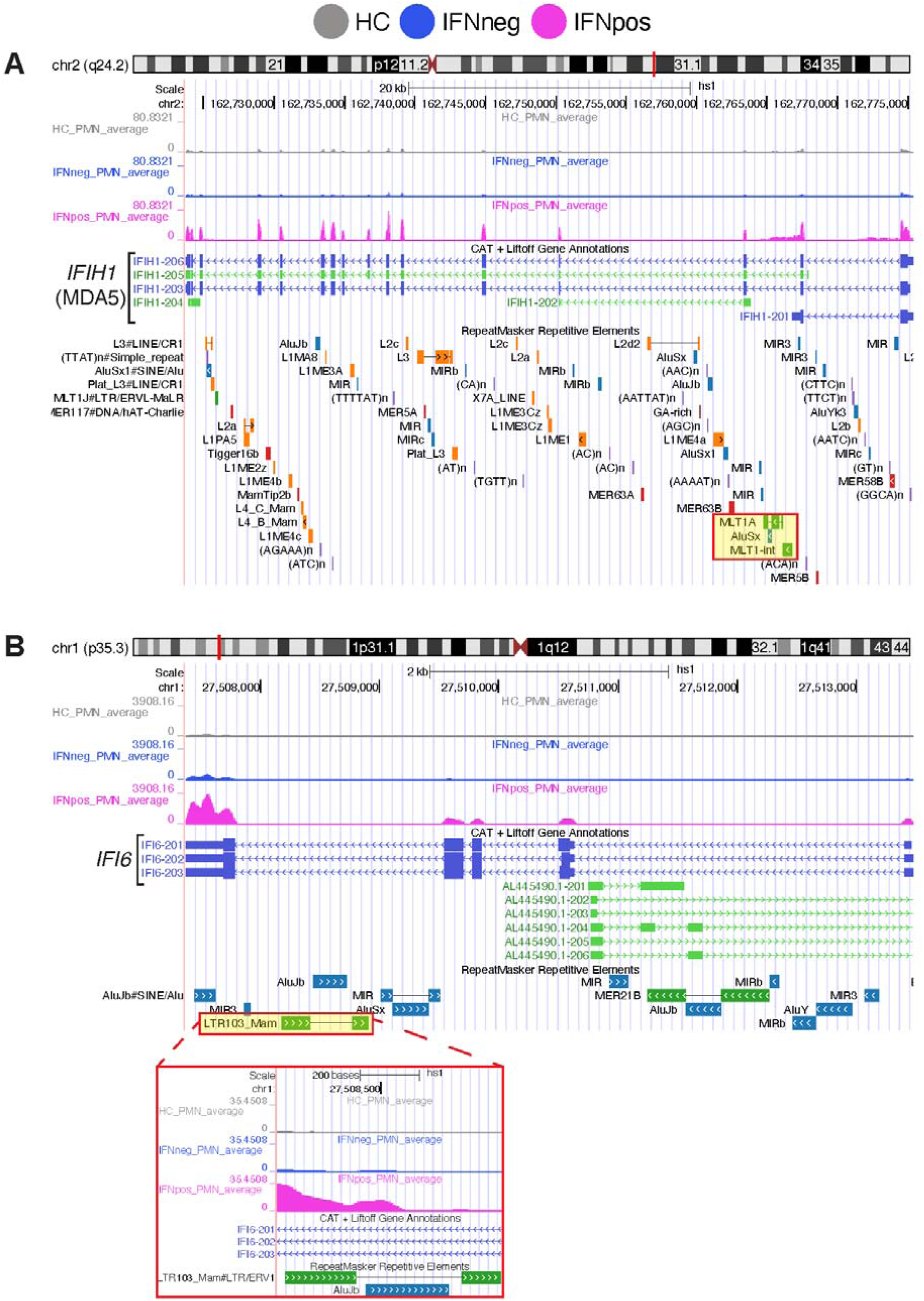
Genome browser views of the intronic MLT1A and MLT1-int elements in IFIH1 locus and intronic LTR103_Mam element in the *IFI6* locus. A, Genome browser view of the *IFIH1* locus encoding MDA5, showing the localization of one of the highest-expressing units of the MLT1A transcription unit and one of the top-expressing units of the MLT1-int family in *IFIH1* intron 2. Note the selective upregulation of transcription in the intron containing this high-expressed TE and the occurrence of annotated splice variants in the flanking exons. B, As in A, but showing the localization of one of LTR103_Mam:ERV1’s highest expressing units in an intron of *IFI6*.

**Supplementary Figure 9.**
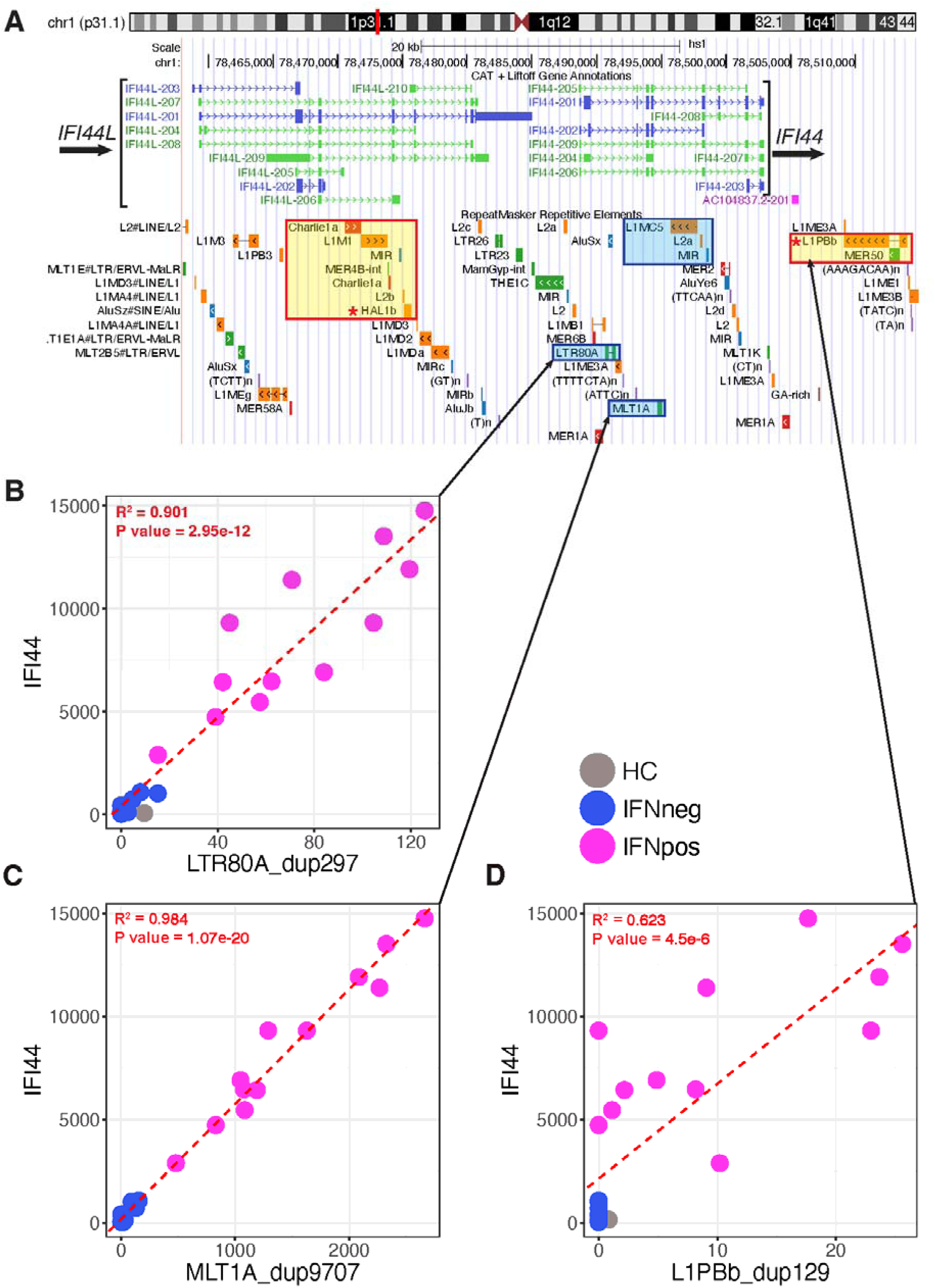
Expanded view of the intronic HAL1B element and LTR80A elements in the IFI44L-IFI44 locus. **A**, Genome browser view of the IFI44L-IFI44 locus. In order, the tracks depict chromosomal location, the annotated splice isoforms of the IFI44L and IFI44 genes, and all repeat/transposable elements in the IFI44L-IFI44 locus. **B**, Scatter plot showing the expression of IFI44 and LTR80A_dup297 of all samples, correlation results denoted in red font which come from the correlation analysis comparing the expression of the gene and the TE in all SLE samples. **C**, Scatter plot showing the expression of IFI44 and MLT1A_dup9707 of all samples, correlation results denoted in red font which come from the correlation analysis comparing the expression of the gene and the TE in all SLE samples. **D**, Scatter plot showing the expression of IFI44 and L1Bb_dup129 of all samples, correlation results denoted in red font which come from the correlation analysis comparing the expression of the gene and the TE in all SLE samples.

